# An RNA interference (RNAi) toolkit and its utility for functional genetic analysis of *Leishmania (Viannia)*

**DOI:** 10.1101/2022.11.14.516516

**Authors:** Lon-Fye Lye, Katherine L. Owens, Soojin Jang, Joseph E. Marcus, Erin A. Brettmann, Stephen M. Beverley

## Abstract

RNA interference (RNAi) is a powerful tool whose efficacy against a broad range of targets enables functional genetic tests individually or systematically. However, the RNAi pathway has been lost in evolution by a variety of eukaryotes including most *Leishmania* sp. RNAi was retained in species of the *Leishmania* subgenus *Viannia*, and here we describe the development, optimization, and application of RNAi tools to the study of *L. (Viannia) braziliensis*. We developed vectors facilitating generation of long-hairpin or “stem-loop” (StL) RNAi knockdown constructs, using Gateway^™^ site-specific recombinase technology. A survey of applications of RNAi in *L. braziliensis* included genes interspersed within multigene tandem arrays such as *QDPR*, a potential target or modulator of antifolate sensitivity. Other tests include genes involved in cell differentiation and amastigote proliferation (*A600*), and essential genes of the intraflagellar transport (*IFT*) pathway. We tested a range of stem lengths targeting the *L. braziliensis* hypoxanthine-guanine phosphoribosyltransferase (*HGPRT*) and reporter firefly luciferase (*LUC*) genes and found that the efficacy of RNAi increased with stem length, and fell off greatly below about 128 nt. We used the StL length dependency to establish a useful ‘hypomorphic’ approach not possible with other gene ablation strategies, with shorter *IFT140* stems yielding viable cells with compromised flagellar morphology. We showed that co-selection for RNAi against adenine phosphoryl transferase *(APRT1)* using 4-aminopyrazolpyrimidine (APP) could increase the efficacy of RNAi against reporter constructs, a useful tool that may facilitate improvements in future work. Thus, for many genes, RNAi provides a useful tool for studying *Leishmania* gene function with some unique advantages.

## 1. Introduction

More than 1.7 billion people are at risk for the ‘neglected tropical disease’ leishmaniasis, with nearly 12 million exhibiting symptomatic disease and more than 50,000 deaths annually, and upwards of 100 million harboring asymptomatic infections [1–5]. *Leishmania* sp. have two distinct growth stages, the promastigote in the sand fly vector, and the intracellular amastigote residing within cellular endocytic pathways in the mammalian host. While the promastigote stage is readily cultured in the laboratory and amenable to molecular techniques, amastigotes require the use of macrophage infection systems or infections of animal models replicating key aspects of human disease.

Different species of *Leishmania* tend to associate with different clinical presentations ranging from localized mild cutaneous disease to more severe visceral or mucosal disease, with host factors also playing key roles [3,6]. *Leishmania* classified within the subgenus *Viannia* are widespread parasites of mammals within South and Central America [7,8]. *L. Viannia* sp. represent one of earliest diverging groups of *Leishmania*, with numerous differences from later-diverging subgenera including development within the insect hindgut, retention of the RNA interference (RNAi) pathway, and often the presence of RNA viruses [7,9,10]. Most importantly this extends to host responses and pathology, especially mucocutaneous disease presentations which most commonly arises from *L. braziliensis* infections and are uncommon in infections by species outside of *Viannia* [11].

In this work we focus on experimental application RNAi pathway, an evolutionarily conserved post-transcriptional gene silencing mechanism in eukaryotes [12]. *Trypanosoma brucei,* a kinetoplastid parasite closely related to *Leishmania,* was the first trypanosomatid and indeed one of the first eukaryotes found to have a functional RNAi pathway [13]. As in other organisms, there RNAi quickly became a key tool for diverse functional genetic analysis, ranging from individual to genome-wide applications [14–16]. In contrast, early studies showed that most species of the related trypanosomatid parasite *Leishmania* lack this pathway, in common with a disjunct group of other eukaryotic microbes [17,18]. These observations have raised many questions about the forces operating on microbes to retain or lose this otherwise universally conserved eukaryotic pathway, such as retention or loss of RNA viruses or retrotransposons [10,19].

While initially disappointing for application towards many *Leishmania* sp., phylogenetic mapping showed that RNAi had been retained in the lineage leading to the subgenus *Viannia*, before its loss in the lineages leading to the remaining species of ‘higher’ *Leishmania* such as *L. major*, *L. donovani* or *L. mexicana* [10]. The discovery of an active RNAi pathway in *Viannia* species raised the possibility that RNAi could be used as useful tool for genetic manipulation, as in other eukaryotes. In *Leishmania* classic gene knockouts by homologous recombination work efficiently, but are challenged by the need to disrupt at least two alleles, and/or the presence of multi-copy genes (although this is rapidly evolving with the implementation of CRISPR/Cas9 based tools [20]). The potential utility of RNAi in *L. braziliensis* emerged in several studies showing the impact of RNAi on cellular protein or RNA levels [10,21].

In this study we provide further evidence for the utility of RNAi in *L. braziliensis*, surveying its activity against a spectrum of gene targets relevant to *Leishmania* biology or chemotherapy, such as flagellar, stage specific or metabolic genes. To facilitate these studies, we applied vectors facilitating the one step production of the dsRNA trigger hairpin-generating “stem-loop” (StL) constructs, using Gateway^™^ (Invitrogen) technology in a single step. A similar approach was described previously in African trypanosomes [22]. We explored several parameters relevant to the efficacy of RNAi, which in turn informed methods to generate hypomorphic loss of function mutations of otherwise lethal knockdowns, and co-selections for elevated RNAi efficacy by negative selection.

## 2. Materials and Methods

### 2.1. *Leishmania* strains and parasite culture

*Leishmania braziliensis* (*Lbr*) M2903 (MHOM/BR/75/M2903) was obtained from D. McMathon-Pratt (Yale School of Public Health) and grown as promastigotes in Schneider’s Insect Medium (Sigma-Aldrich cat. No. S9895) supplemented with 10% heat-inactivated fetal bovine serum (FBS), 2 mM L-glutamine, 500 units 500 units ml^−1^ penicillin and 50 μg ml^−1^ streptomycin (Gibco No. 5070). *Leishmania braziliensis* M2903 SA (MHOM/BR/75/M2903) SA was obtained from S.C. Alfieri (Universidade de São Paulo, Brasil). The SA line had been adapted for growth and was serially maintained as amastigotes at 34°C in modified UM-54 medium [23].

### 2.2. *Leishmania* transfection

Stable transfection of *L. braziliensis* M2903 strain (MHOM/BR/75/M2903) was performed using the high-voltage procedure [24]. Parasites were grown to mid-log phase, pelleted at 1300 *g*, washed once with cytomix electroporation buffer (120 mM KCl, 0.15 mM CaCl_2_, 10 mM K_2_HPO_4_, 25 mM HEPES-KOH, pH 7.6, 2 mM EDTA and 5 mM MgCl_2_) and resuspended in cytomix at a final concentration of 1 × 10^8^ cells ml^−1^. For transfection, 10 μg DNA was mixed with 500 μl of cells and electroporated twice in a 0.4 cm gap cuvette at 25 μF, 1400 V (3.75 kV cm^−1^), waiting 10 sec between electroporations. Cells were then incubated at 26° C for 24 hours in drug-free media and then plated on semisolid media containing the appropriate drug to select clonal lines. For selections using blasticidin deaminase (*BSD* gene), hygromycin phosphotransferase (*HYG* gene), streptothricin acetyltransferase (*SAT* gene) and the bleomycin-binding protein from *Streptoalloteichus hindustanus* (*PHLEO* gene) markers, parasites were plated on 10-20 μg ml^−1^ blasticidin, 30-80 μg ml^−1^ hygromycin B, 50-100 μg ml^−1^ nourseothricin and 0.2-2 μg ml^−1^ phleomycin, respectively (ranges reflect differences when using drugs singly or in combination). Colonies normally appeared by 14 days, at which point they were recovered, grown to stationary phase in 1 ml and passaged with the appropriate drugs. Plating efficiencies range from 60% to 95% for the un-transfected *L. braziliensis* M2903 strain; transfection efficiencies varied from 2 to 50 colonies per μg of *Swa*I-digested pIR vector controls.

### 2.3. Western-blot analysis

*Leishmania* promastigotes were collected and resuspended in phosphate buffered saline (PBS) at 1×10^8^ cells/ml. Cell extracts were prepared and Western blots performed after separation by SDS-polyacrylamide gel electrophoresis as previously described [25]. For HGPRT, primary antibodies were anti-*L. donovani* HGPRT and APRT antiserum [26] was used at a titer of 1:5000 and 1:1000 respectively. For normalization anti-*L. major* H2A [27] was used at a titer of 1:100,000, with goat anti-rabbit IgG as the secondary antibody (1:20,000, Licor Inc.). Similar procedures were used for QDPR western blot analysis with extracts from 1.6 × 10^7^ cells. Gels were transferred to nitrocellulose membranes, which were blocked with a 5% skim milk solution and incubated with 1:500 dilution of rat anti-QDPR[28], a 1:1000 dilution of rabbit anti-PTR1 [29] or anti-*L. major* H2A as described above. IRDye™ anti-rat or rabbit goat immune globulin G were used as the secondary antisera at 1:10,000 dilution. Antibody binding to blots was detected and quantified using an Odyssey infra-red imaging system (Li-Cor).

### 2.4. *Quininoid* dihydropteridine reductase assay

Parasites were harvested at log phase (4-6 × 10^6^ cells/ml) and collected by centrifugation at 1,250 *g* for 10 min at 26° C, washed twice with PBS, and resuspended at 2 × 10^9^ cells ml^−1^ in 10 ml of Tris-Cl, pH 7.0, with 1 mM EDTA and a mixture of protease inhibitors as described [30]. Cells were lysed by three rounds of freeze thawing and sonication, and the extracts clarified by centrifugation at 15,000 x *g* for 30 min at 4°C. Protein concentrations were determined using Qubit Fluorometric Quantification (Invitrogen). *Quininoid* dihydropteridine reductase activity was measured at 25°C as described [31] using *quinonoid* dihydrobiopterin generated continuously by horseradish peroxidase mediated oxidation of H_4_-biopterin. The standard reaction mixture contained 50 mM Tris-HCl, pH 7.2, 20 μg of horseradish peroxidase, 0.1 mM H_2_O_2_, 20 μM of H_4_-biopterin, 100 μM NADH, and purified QDPR or parasite lysates; all components were incubated for 3 min prior to initiation of reaction by addition of H_4_B. The activity was measured by monitoring NADH consumption at 340 nm (ε_340_ for NADH is 6200 M^−1^ Cm^−1^) in a Beckman DU-640 spectrophotometer.

### 2.5. Luciferase assay

Logarithmic growth phase promastigotes (10^6^) were suspended in 200 μl media containing 30 μg/ ml of luciferin (Biosynth AG) and added to a 96-well plate (Black plate, Corning Incorporated, NY, U.S.A.). The plate was imaged after 10 min using a Xenogen IVIS photoimager (Caliper LifeSciences), and luciferase activity quantitated as photons/sec (p/s).

### 2.6 Transmission electron microscopy

*Leishmania* promastigotes were harvested in logarithmic growth phase and fixed in 2% paraformaldehyde/2.5% glutaraldehyde (Polysciences Inc., Warrington, PA) in 100 mM phosphate buffer, pH 7.2, for 1 hr at room temperature. Samples were washed in phosphate buffer and postfixed in 1% osmium tetroxide (Polysciences Inc., Warrington, PA) for 1 hour, rinsed extensively in water, and block stained with 1% aqueous uranyl acetate (Ted Pella Inc., Redding, CA) for 1 hr. Following several rinses in water, samples were dehydrated in a graded series of ethanol solutions and embedded in Eponate 12 resin (Ted Pella Inc.). Sections of 95 nm were cut with a Leica Ultracut UCT ultramicrotome (Leica Microsystems Inc., Bannockburn, IL), stained with uranyl acetate and lead citrate, and viewed on a JEOL 1200 EX transmission electron microscope (JEOL USA Inc., Peabody, MA).

### 2.7. Construction of the integrating pIR-GW destination vectors facilitating generation of StL constructs using Gateway site specific recombinase

We first assembled a construct bearing a PEX11-MYC ‘loop’ fragment flanked by inverted Gateway® cassettes; these contain *ccdB*, a lethal gene that targets DNA gyrase, and *CmR* encoding chloramphenicol-resistance, flanked by two *attR* sequences necessary for site-specific recombination (*attR1-ccdB-CmR-attR2* ; www.Invitrogen.com), For propagation, all constructs bearing the Gateway cassettes were propagated in *E. coli* DB3.1 which contains a *gyrA462* mutation conferring resistance to *ccdB* toxicity. First, the PEX11-MYC loop was excised from pIR1*SAT*-GFP65-StL (B4733) [7,9,10] and inserted into pGEMT yielding pGEMT-stuffer (B5974). The Gateway cassette (*attR1-ccdB-CmR-attR2*) was amplified from the pDONR221 (Invitrogen) with primers S1 and S2, and inserted by blunt end ligation into the *Nhe*I site of pGEMT-stuffer (B5974), yielding pGEMT-stuffer -one GW (B6158). The second GW cassette was inserted by blunt end ligation into the pGEMT-stuffer -one GW *Avr*II site, in inverted orientation to the first cassette (with both in divergent orientation relative to the *ccdB/CmR* ORFs), yielding (B6218).

Plasmid pGEMT-stuffer - 2 GW divergent contains a 3877 bp *SphI/HindIII* fragment bearing the inverted Gateway cassettes and loop. This fragment was blunt-end ligated into the *Sma*I site of pIR1*SAT* (B3451), yielding pIR1*SAT*-GW (B6223) (**Supplemental Table S1**). Similarly, the inverted Gateway/loop fragment inserted into the *Bgl*II (B) site of various pIR vectors by blunt end ligation, yielding the final vectors pIR1*SAT*-GW (B6223), pIR1*HYG*-GW (B6544), pIR1*PAC*-GW (B6543), pIR1*BSD*-GW (B6542) pIR2*HYG*-GW (B6563) (**Supplemental Table S1**). The sequence of pIR1*HYG*-GW is provided in **Supplemental File 1**.

### 2.8. Generation of target stem-loop (StL) constructs for RNAi

The molecular constructs used in this work and their synthesis are summarized in **Table S2**. Briefly, for StL constructs the ‘stem’ was obtained by PCR, inserted into the pCR8/GW/TOPO vector by TA cloning, (Invitrogen # K250020), and their orientation (same direction as *attL2*) confirmed by sequencing and restriction digestion. The stems were transferred from the pCR8/GW/TOPO donor vector to the pIR1-GW or pIR2-GW destination vectors described above; these contain sequences from the parasite small subunit rRNA locus to enable integration into the genome, and inverted *attR1-ccdB-CmR-attR2* cassettes. The gene of interest in pCR8/GW/TOPO was transferred to the pIR-GW vector with LR Clonase II (Thermo Fisher) in an overnight reaction at room temperature. Reactions were terminated by incubating with proteinase K for 1 h at 37 °C (Gateway Technology with Clonase ^™^ II manufacturer’ protocol, Version A: 24 June 2004). Reactions were transformed into *E.coli* TOP10 ^™^ or DH5α (which select against both *ccdB* cassettes). All final stem-loop (StL) constructs named as StL expressers were confirmed by restriction enzyme digestion and DNA sequencing (**Figure 1**). Prior to transfection, constructs were digested with *Swa*I to expose the SSU rRNA segments mediating homologous integration.

**Figure 1.**
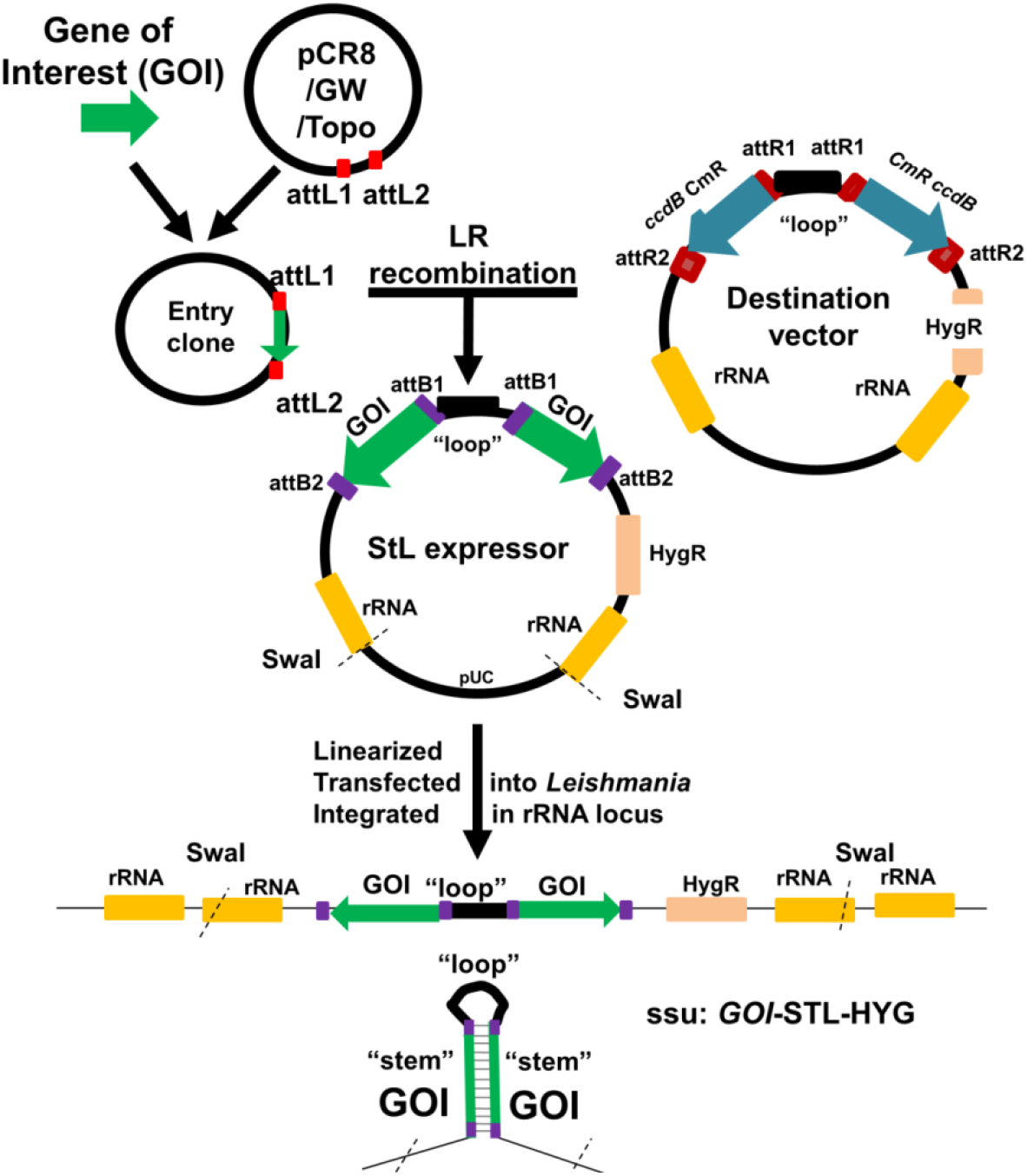
Flowchart for generation of entry and StL constructs using Gateway site-specific recombinase technology followed by introduction into *Leishmania* to express dsRNA. Each stem for GOI (gene of interest) was amplified by PCR then cloned into pCR8/GW/TOPO vector (Invitrogen) to generate an “entry” clone. This pCR8/GW/TOPO vector includes an *attL*1 and *attL*2 sites for recombination-based transfer of the gene of interest into the desired pIR-GW destination vector, which bears to opposing *ccdB/CmR* cassettes each flanked by *attR*1 and *attR*2 sites. Following site-specific recombination by the Gateway LR reaction, the desired StL for each GOI was obtained and confirmed. For biological tests, each StL DNA was linearized with *Swa*I, electroporated into *Leishmania*, where it integrated into the small subunit ribosomal RNA locus (SSU). There it is transcribed by the strong rRNA pol I promoter to generate RNAs whose dsRNA regions are processed by the RNAi machinery.

## 3. Results

### 3.1. Rapid generation of ‘stem-loop’ constructs as RNAi triggers

To trigger the RNAi response, dsRNA generating ‘stem-loop’ constructs have been used in previous studies, often assembled in three steps with two ‘stem’ segments cloned in opposite orientations, separated by a short spacer/loop [10]. To accelerate this process, we utilized Gateway^™^ (Invitrogen) technology, incorporating precise, site-specific recombination [32].[22]. First we engineered a ‘destination’ expression vector, based on the *Leishmania* pIR vector series which achieves high levels of RNA expression following integration into the ribosomal small subunit RNA locus [10,33]. The final destination vector bore two Gateway recipient cassettes, arranged in inverted orientation and separated by a short segment destined to become the ‘loop’ inserted into one of the strong expression sites of pIR vectors (**Figure 1**). Each cassette contained a positive *CmR* and negative *ccdB* marker, flanked by *attR*1 and *attR*2 sites (pIR-GW vector). Each ‘stem’ to be tested was inserted into an ‘entry’ vector (pCR8/GW/TOPO, flanked by donor *attL*1 and *attL*2 sites (**Figure 1**)). Gateway Reactions bearing the donor vector and entry DNAs were performed *in vitro*, and transformed into *E.coli* TOP10 or DH5α^™^ which selected against the presence of both target *ccdB/CmR* cassettes, which was confirmed by loss of chloramphenicol resistance. Typically we obtained numerous recombinants of which more than 70% bore the expected configuration of the desired StLconfiguration, and one correct representative was selected for introduction into *Leishmania*. Prior to transfection, constructs were digested with *Swa*I to expose the SSU rRNA termini to direct integration into the SSU rRNA locus, where strong expression is driven from the rRNA promoter (**Figure 1**).

### 3.2. Testing the effect of stem length on RNAi activity using *L. braziliensis HGPRT*-StL and luciferase-StL series constructs

In *T. brucei*, stem lengths ranging in length between 100 and 1500 bp are effective in knocking down target genes [34,35], and stem lengths as short as 29-100 nt are active in other metazoan species [36,37]. To explore this in *Leishmania*, we varied the stem length in RNAi constructs targeting an integrated firefly luciferase reporter gene (*LUC*) as well as an endogenous cellular gene (*HGPRT*), assayed following transfection into WT *Lbr*. For *HGPRT* increasing the stem from 494 nt to 1005 nt reduced expression from 55% to 92 % (**Figure 2A**). No strong differences were seen between similarly sized stems targeting the *HGPRT* ORF or 3’ untranslated region (**Figure 2A**).

**Figure 2.**
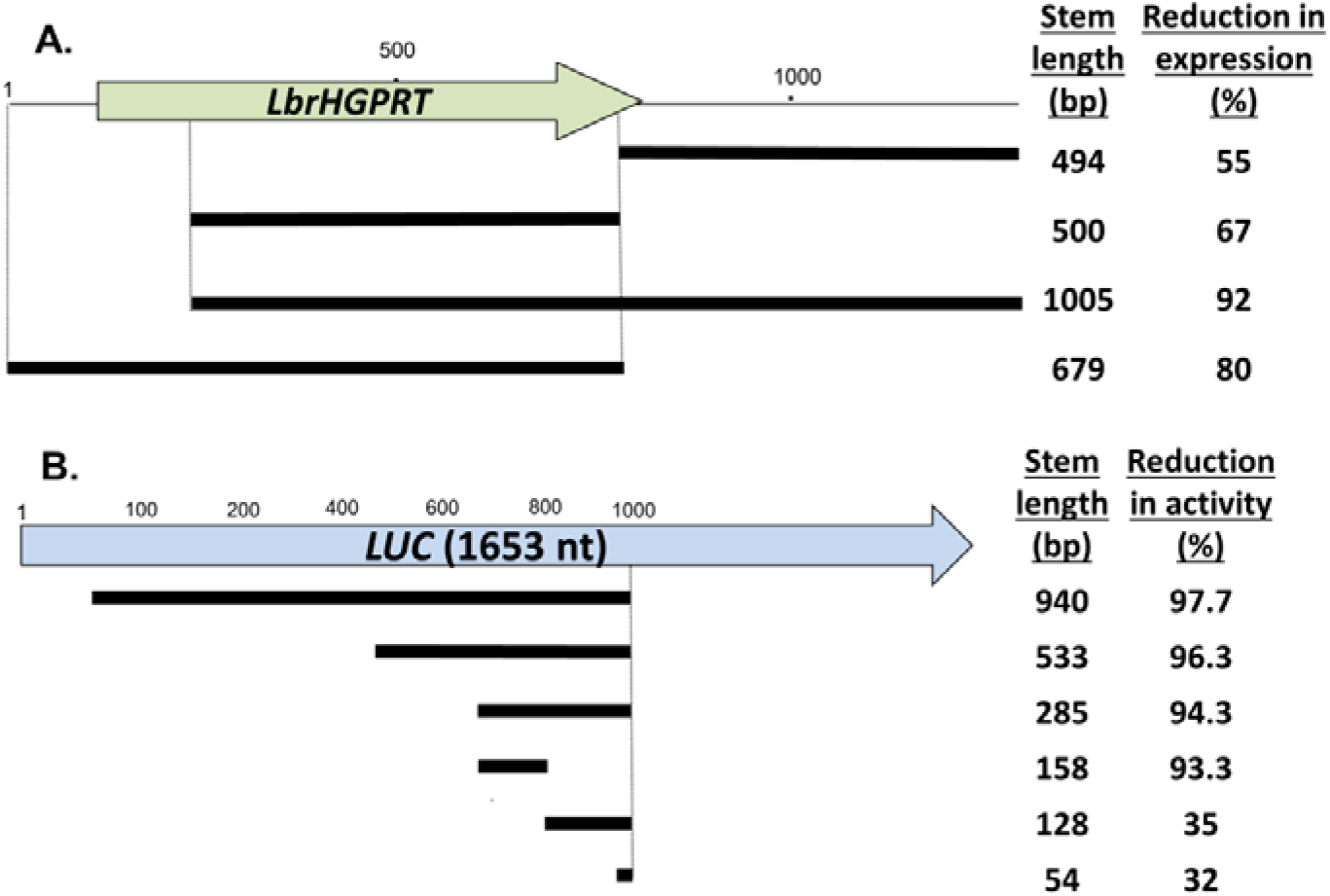
The effect of stem length on RNAi activity. **(A)** Targeting the endogenous *HGPRT*. Various stem length *HGPRT* StL constructs were transfected into *L. braziliensis,* and HGPRT expression assessed in two colonies by western blot with anti-HGPRT antisera and anti-H2A antisera for normalization (**Supplementary Figure S1**). The percent expression relative to WT *Lbr* is shown. **(B)** Targeting an introduced luciferase reporter. Various stem length *LUC* StL constructs were transfected into an *Lbr* line expressing *LUC*. Luciferase activity was measured and the percent reduction in activity calculated relative to the untransfected parental Lbr-LUC-expressing line.

For luciferase, we tested *LUC* StL constructs following introduction into a *Lbr*M2903 transfectant stably expressing high levels of luciferase activity (**Figure 2B**). As with *HGPRT*, the longer luciferase stems resulted in greater reductions in *LUC* activity (**Figure 2B**). While stems of 158 bp or greater showed strong reductions (>93 %), stems of 128 or 54 nt showed much smaller effects (35-32 %; **Figure 2B**). Preliminary analysis did not suggest a strong association with siRNA abundance ‘hot spots’ [19] nor synergy when two weak constructs were transfected simultaneously (not shown). These data suggested differences in the dsRNA trigger length dependency in *Leishmania* relative to trypanosomes or other organisms, although further studies will be required to explore its mechanism or generality. These data suggested that for the strongest effect stem lengths of >500 nt were preferable, with a significant dropoff below 128 nt.

### 3.3. StL-mediated specific RNAi of the metabolic target *quinonoid* dihydropteridine reductase (*QDPR*) interspersed within a tandem repeatedly gene array

We examined the efficacy of StL RNAi against the *L.braziliensis quinonoid* dihydropteridine reductase (*QDPR*), a component of the reduced pteridine pathways playing key roles in *Leishmania* [28]. In all *Leishmania, QDPR* genes are interspersed in a tandem array with two other genes, *ORFq* (*q*: a hypothetical protein) and *β7-proteasome* (*β7*: 20S proteasome β7 subunit), the latter gene being essential where tested in other species (**Figure 3A**; [28]). This rendered selective deletion of the interspersed *QDPR* targets considerably more challenging, but provided a suitable opportunity for testing the utility of RNAi in this context.

**Figure 3.**
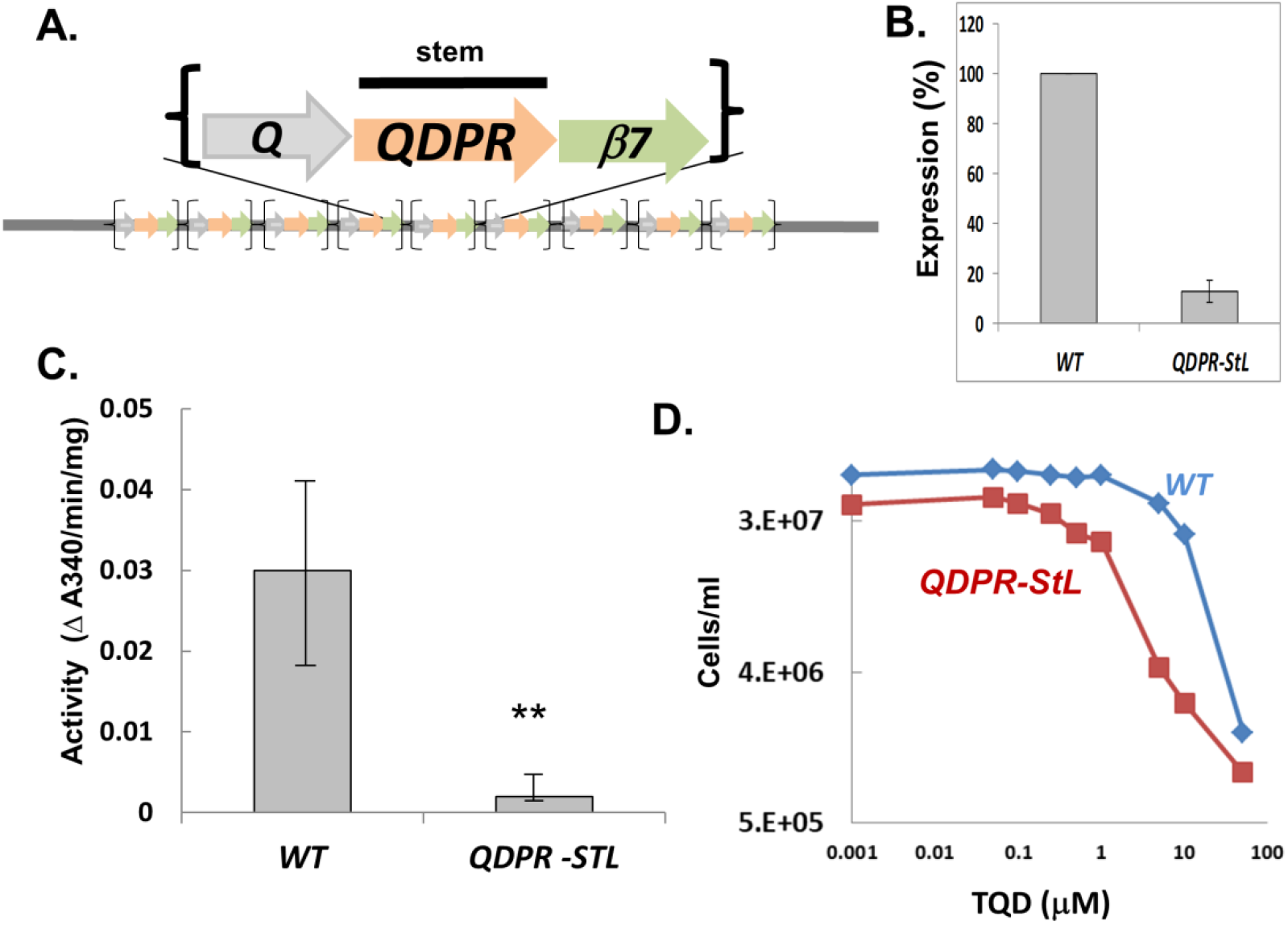
RNAi of the tandemly dispersed gene *QDPR*. **(A) *QDPR* gene organization in *L. braziliensis***. *QDPR* genes are tandemly repeated with *ORFq* (*Q*) and *β7 proteasome (β7)* up to 9 times in *L. braziliensis* genomes. Proteasome subunits are known to be essential, while the requirement for *ORFq* has not been tested. **(B) QDPR protein expression in *QDPR*-StL knockdowns.** QDPR was detected with anti-QDPR antiserum and expression was calculated relative to that of H2A, from western blots as described in the methods. **(C) QDPR enzymatic activity in *QDPR*-StL knockdowns.** QDPR enzymatic activity was assayed and normalized to total cellular protein. Data were from three independent experiments each performed in triplicate. ** indicates *p* < 0.0235. **(D) Growth inhibition by inhibitor TQD.** WT and *QDPR*-StL transfected *Lbr* were inoculated into media containing 0–50 μM 5, 6, 7, 8-Tetrahydro- 2, 4-quinazolinediamine (TQD) at 2 × 10^5^ cells/ml and allowed to grow until WT had reached late log phase, at which time parasite numbers were determined. The experiment was repeated three times, with results similar to that shown. WT *Lbr* is shown in blue, *QDPR*-StL in red.

We introduced a StL construct bearing a 588 nt *QDPR* stem into WT parasites successfully. Relative to WT, QDPR protein was reduced more than 87.4% **(Figure 3B)** while QDPR activity was reduced by more than 88% individually in StL knockdowns (**Figure 3C**). The *Lbr QDPR* StL knockdowns grew normally in culture, suggesting that RNAi specifically targeted *QDPR* without significantly impacting the flanking essential proteasome subunit. Thus RNAi can be used to successful probe the consequences of metabolic gene expression depletion in this most challenging context.

One functional consequence of *QDPR* ablation was shown by the increased sensitivity of the *QDPR* StL knockdown to the antifolate compound TQD (5, 6, 7, 8-Tetrahydro- 2, 4-quinazolinediamine) (**Figure 3D**) [38]. The *QDPR* knockdown parasite was 20 fold more sensitive, with an EC50 of 0.15±0.015 vs 3.49±0.59 μM for WT wild type. These data support the utility of RNAi knockdowns in *Lbr* as as probes of pteridine metabolism and drug action.

### 3.4. RNAi of an *L. braziliensis* gene important for amastigote replication

An important area of *Leishmania* biology is the study of genes that impact survival of the amastigote stage in the vertebrate host. Many (not all) *Leishmania* species can differentiate to amastigote-like forms in culture (axenic amastigotes), when grown at conditions resembling those within the parasitophorous vacuole, eg elevated temperature and low pH [39,40]. Here we made use of a clonal derivative of *Lbr* M2903 (LbrM2903SA2) which had been adapted for growth as axenic amastigotes [23].

As a test we focused on the *A600* locus, which in *L. mexicana* comprises four genes whose deletion had little impact on promastigotes, but precluded axenic amastigote replication *in vitro* [41]. The *A600* copy number varies amongst species [41], with only two found in *Lbr* (*A600-1* and *A600-4*) which show 88% nucleotide identity. We targeted these sequences simultaneously by a single StL construct using the *A600-1* sequence, which shows long stretches of identity relative to *A600-4* (118 nt, 51 nt and 45 nt).

We transfected the *LbrA600*-StL construct into *Lbr* SA2 promastigotes, obtaining many clonal transfectants all of which grew normally. Several were then inoculated into axenic amastigote growth medium, where they showed a severe growth defect, with 10-fold fewer amastigotes than WT (**Figure 4**), similar to the results obtained with *L. mexciana A600* knockouts. These results establish the utility of RNAi knockdowns for the study *L. braziliensis* genes involved in amastigote differentiation and proliferation.

**Figure 4.**
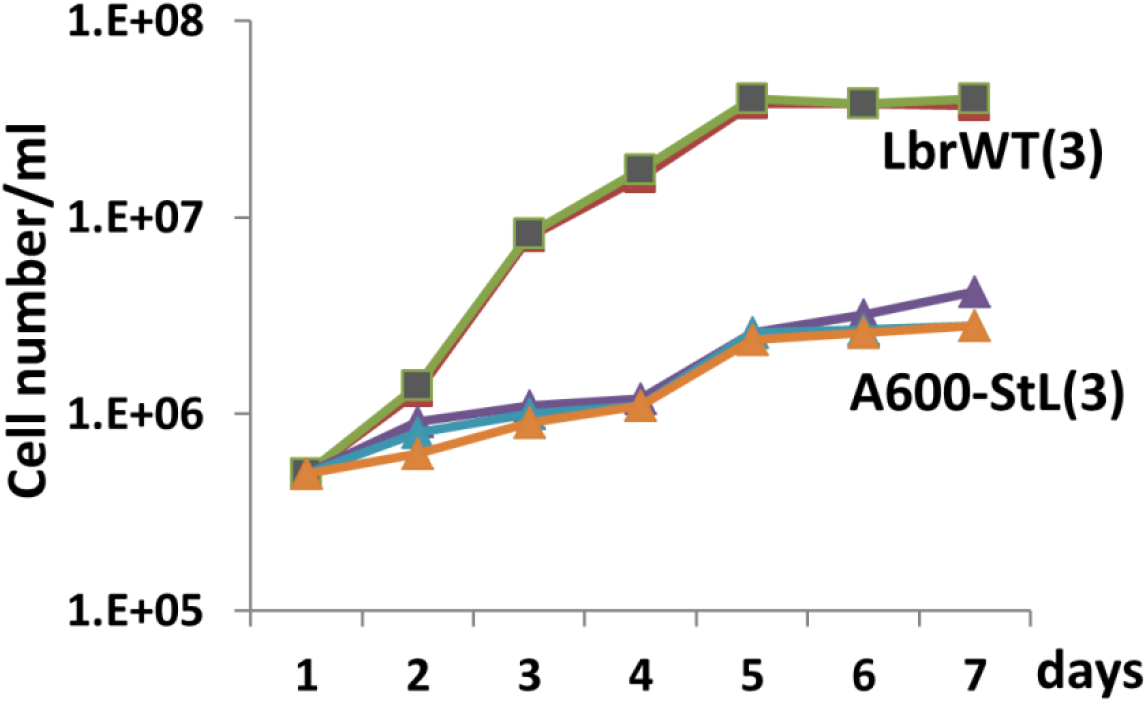
Impact of *LbrA600*-StL amastigote growth in *L. braziliensis*. WT or *A600-*StL transfectants grew normally as promastigotes; the figure shows parasites inoculated at a density of 5×10^5^ /ml and the cell numbers determined daily for 1 week. WT (■) was repeated three times along with 3 different *A600*-StL knockdown clonal lines (#1-3; ▲)

### 3.5. Targeting essential genes of the *L. braziliensis* intraflagellar transport (IFT) pathway

The *Leishmania* flagellum plays key roles in the promastigote and amastigote stages, and previously we reported successful knockdowns of the paraflagellar rod proteins PFR1 and PFR2 which showed phenotypes comparable to complete deletions [10]. We extended these studies to a set of genes associated with intraflagellar transport (IFT), which mediates transport of cargo in both anterograde and retrograde directions and is required for proper flagellar assembly [42].

StL constructs targeting *Lbr IFT122, IFT140, IFT172,* representing both the anterograde and retrograde IFT, were transfected into WT *Lbr* (Table 1). However, we were unable to recover transfectant colonies, despite multiple attempts and success targeting a nonessential gene, *LbrPFR1*-StL (**Table 1**). To establish that this result was dependent on the RNAi pathway, we introduced these same constructs into an RNAi-deficient mutant obtained by homozygous deletion of the *AGO1* gene (*Δago1*^−^), an Argonaute family protein encoding the key ‘slicer’ activity required for RNAi activity in *Lbr* [10,43]. Now, all *LbrIFT* StL constructs successfully yielded transfectants (**Table 1**), establishing that RNAi of the StL-derived dsRNA trigger was responsible for the lack of transfectants.

**Table 1.** RNAi of several IFT genes in *Leishmania braziliensis* is lethal.

| STL construct | No. transfectants with WT | No. transfectants with *Δago1*<sup>-</sup> |
| --- | --- | --- |
| *PFR1*-StL | 243-360 | 360 |
| *IFT122*-StL (retrograde) | 0 | 460 |
| *IFT140*-StL (retrograde) | 0 | 265 |
| *IFT172*-StL (anterograde) | 0 | 289 |
| Negative control | 0 | 0 |

These data are consistent with other studies in trypanosomes and *Leishmania* showing that *IFT* gene ablation displays an array of phenotypes, including essentiality [44–47].

The table shows the number of colonies (per plate) obtained after of transfection of various StL constructs into WT or RNAi-deficient *Δago1^−^ Lbr. PFR1*-StL was used as a positive control [10].

### 3.6. Systematic generation of hypomorphic mutants by exploiting the stem length-dependency of RNAi in *L. braziliensis*

While plus/minus tests of gene essentiality are useful they provide little information about the role of the encoded protein within the cell. When inducible systems are available, observations following shutoff and prior to cell death can be informative, however presently this can only be achieved using conditional degradation domains in *Leishmania* [48]. The studies of the stem length-dependency of the efficacy of RNAi described in Section 3.2 above suggested that it could be possible to engineer partial loss of function mutants through successively reducing the stem length, until a point was reached where viable cells could be obtained which ideally might show informative defects. We tested this with the *IFT140* gene, which is essential in trypanosomes and probably *Lbr* (**Table 1)**. Curiously, viable deletion mutants with different phenotypes were possible in *L. major* and *L. donovani* [46,47] suggesting there is considerable variation amongst amongst *Leishmania* species for the consequences of IFT ablation.

Transfection of a 941 nt *IFT140* StL construct yielded no transfectants in WT *Lbr* (**Table 1**), however as the stem length was reduced to 562 nt and further to 131 nt, increasing numbers of transfectants could be recovered (**Figure 5A**). Notably, the 562 nt stem yielded about 10% as many transfectants, and the cells recovered from these colonies grew slowly in culture (**Figure 5A**). In contrast, transfectants recovered with constructs with stems shorter than 562 nt were recovered at normal frequencies, and grew like WT. This suggested that the 562 StL *IFT140* transfectant was a candidate hypomorph. Transmission EM of one clone confirmed this supposition, as the knockdown cells exhibited modified shape and accumulated vesicles in the flagellar pocket similar to the phenotype in trypanosomes in which ablation of *IFT140* is initiated by conditional RNAi prior to cell death [44] (**Figure 5C**). These data suggest that the stem-length dependency of RNAi in *L. braziliensis* will prove a useful tool in generating informative, hypomorphic mutants of otherwise essential genes, facilitating inquiries into their cellular targets and/or mechanism of action.

**Figure 5.**
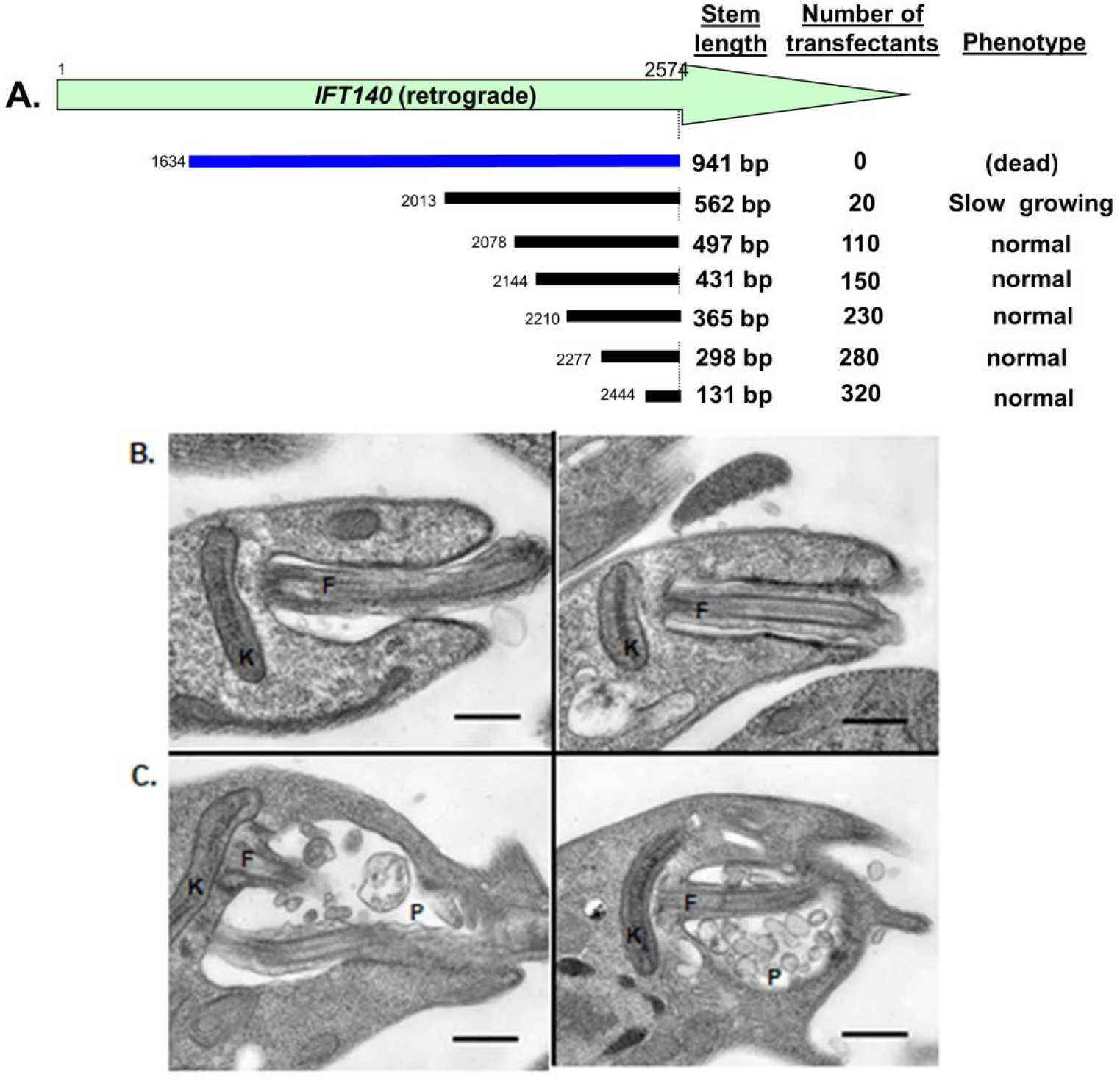
Systematic identification of viable hypomorphic RNAi mutants. **(A)** Map of the *IFT140* stems tested, number of colonies obtained after transfection, and the growth properties (when viable). **(B)** Transmission EM of WT *Lbr* showing normal kinetoplast (K) and flagellum (F). **(C)** Transmission EM of hypomorphic *LbrIFT140*-StL (562 nt stem) cells. Defects in the parasite kinetoplast (K) and flagellum are evident (F) along with accumulation of vesicles within the flagellar pocket (P); not shown is that these cells are more rounded.

### 3.7. A negative selection system for enhancement of RNAi activity in *Leishmania braziliensis*

It is often helpful to have screens or selections to monitor the efficacy of RNAi under various circumstances; for example, in screening for genes acting within the RNAi pathway , or when RNAi exhibits considerable clonal variability, as in some fungi [49–51]. GFP and luciferase based screens are commonly used, which both perform well in *Leishmania* [10], but here we sought a selectable system which could facilitate other applications.

We tested the 4-aminopyrazolpyrimidine/adenine phosphoryl transferse 1 (APP/*APRT1*) system in *L. braziliensis*. APRT converts APP into a toxic metabolite inhibiting *Leishmania* growth, and thus cells with decreasing APRT levels show increasing resistance to APP. Importantly, *APRT1* is not essential in most media to the alternative salvage route through adenosine aminohydrolase [52].

First, we introduced an *APRT1*-StL construct into WT *Lbr*, where transfectants were readily obtained and grew normally. Western blot analysis confirmed significant reduction to undetectable level in APRT1 expression (**Figure 6A**). Unlike WT parasites whose growth was completely inhibited by 500 μM APP, *LbrAPRT1*-StL transfectants were able to grow at the highest concentration tested, with only a small reduction in growth rate at 1 mM APP (**Figure 6B, C**). Thus, RNAi knockdown of APRT1 leads to APP resistance as expected.

**Figure 6.**
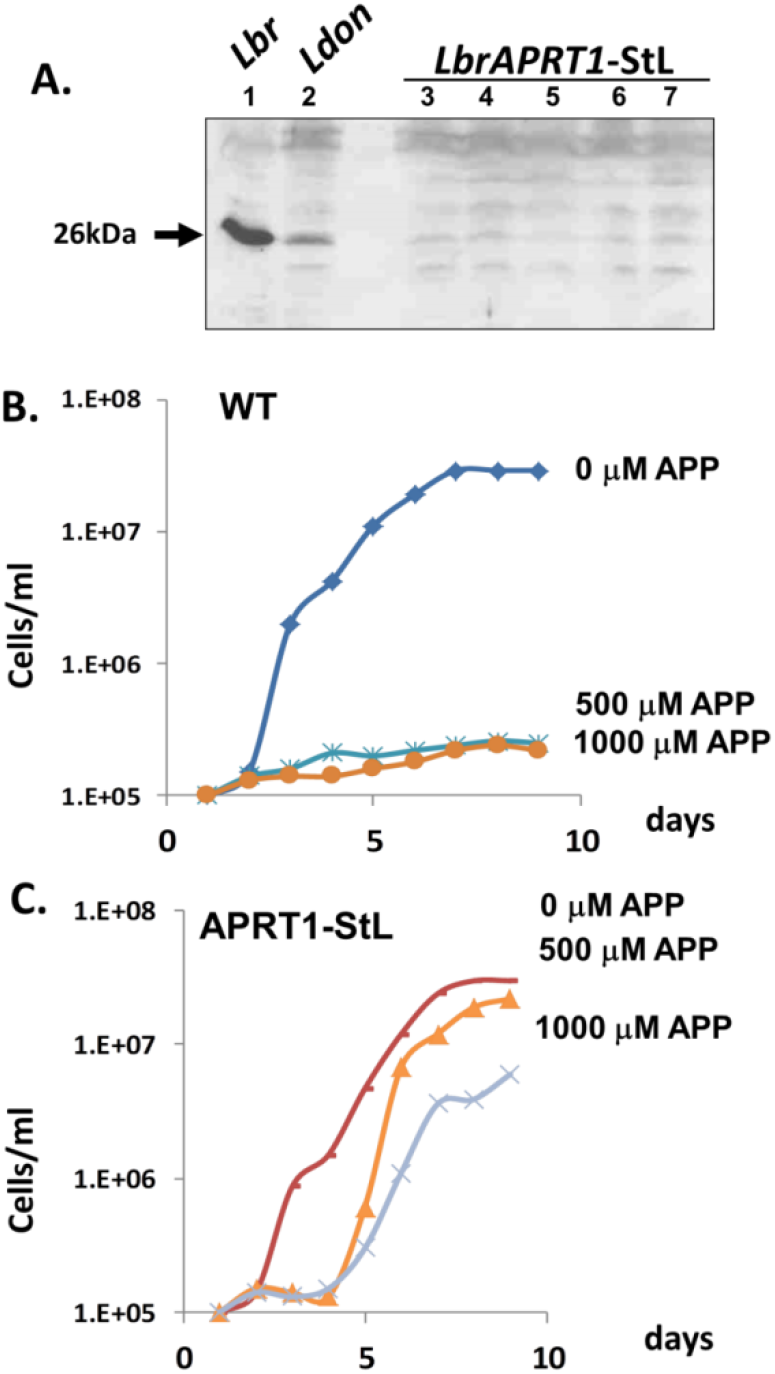
RNAi of the *APRT1* gene yields increased APP resistance. **(A)** Transfection of *APRT1*-StL leads to loss of APRT1 expression. The figure shows a western blot of promastigotes tested against anti-APRT1 antisera. Lane 1. *L. braziliensis* WT; lane 2, *L.donovani* WT; lanes 3-7. *Lbr APRT1*-StL clones 1 to 5. **(B, C)** Growth inhibition by APP in the indicated concentrations is shown. B: WT *Lbr*; C: *Lbr APRT1*-StL knockdown clone 1.

We developed a dual stem-loop reporter parasite that simultaneously expresses an *APRT1*-StL along with a *LUC*-StL and luciferase. This allows manipulations of RNAi through APP inhibition of APRT1 to be tracked through simple luciferase assays. To illustrate its utility, *APRT1-StL/LUC-StL/LUC* parasites were then plated on increasing concentrations of APP, which would select for low APRT1 expression through RNAi. Parasites were grown in 50% of the selective concentration of APP used in plating during subsequent growth and analysis.

In the absence of APP, luciferase expression was reduced 122 fold in the *APRT1-StL/LUC-StL/LUC* parasites (Figure 7), slightly less than the 200- 300 fold reduction seen previously [10]. In the *APRT1-StL/LUC-StL/LUC* parasites subjected to APP selection, luciferase activity decreased further from 2.9 to 6.8 fold, as the APP concentrations were increased from 125 to 1000 μM (**Figure 7**). Overall relative to WT parasites, the efficacy of LUC silencing increased from 122 to nearly 830 fold. Thus, in one step we were readily able to identify parasite showing elevated levels of RNAi. This proof of principle experiment establishes a useful tool for both monitoring and manipulating RNAi activity amongst clones or mutants that may prove helpful in future studies.

**Figure 7.**
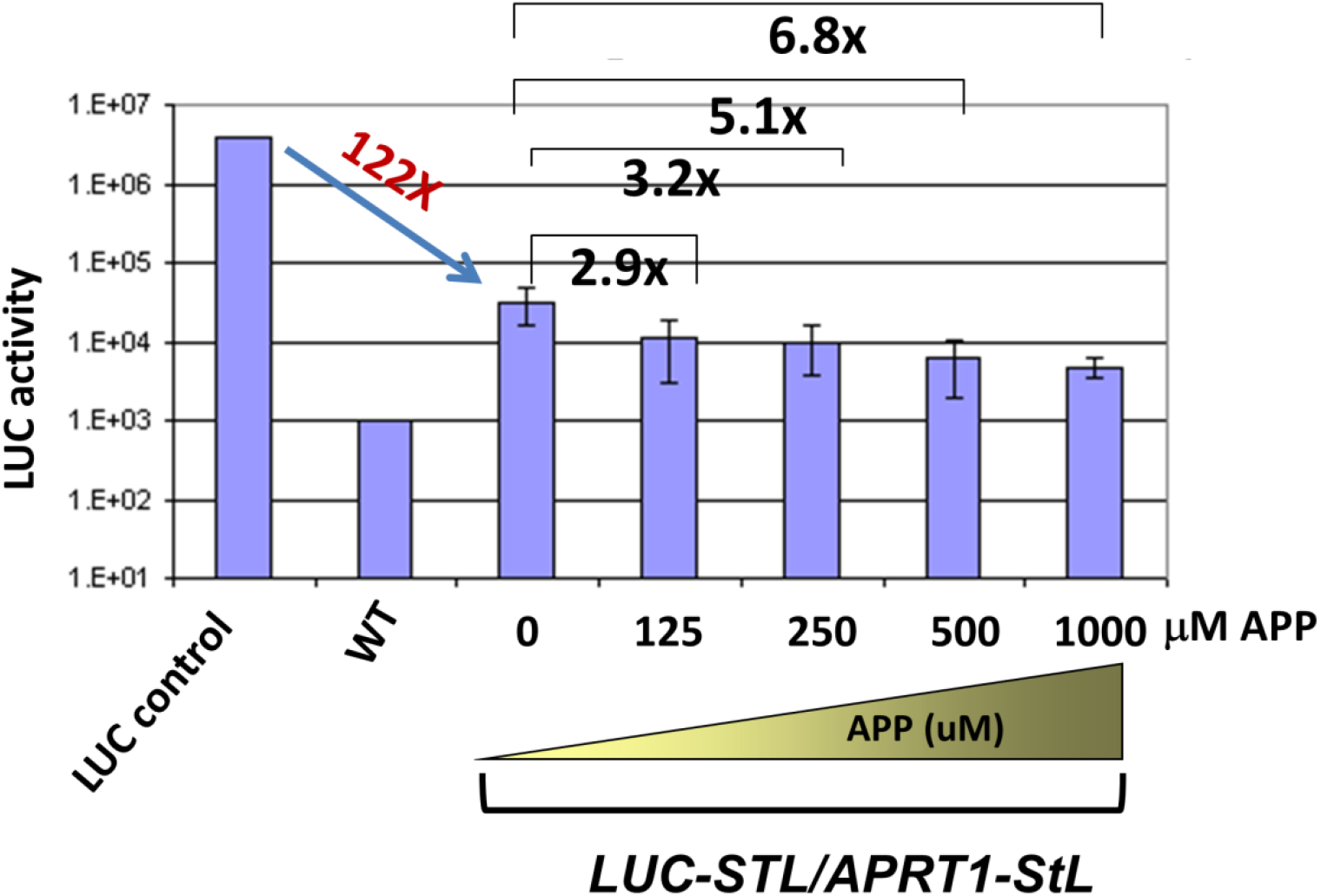
A negative selectable system to identify enhanced RNAi activity in *Leishmania braziliensis*. Double StLs (*LUC* and *APRT1*) expressed in a single construct were transfected in the *Lbr LUC* reporter line and selected on M199 plates containing various concentrations of APP. Colonies were readily obtained and grown in APP, and luciferase activity. The average and standard deviations calculated from at least 10 transfectant colonies is shown.

## 4. Discussion

RNA interference has proven a valuable tool for the study of gene regulation in many eukaryotes including African trypanosomes. While lost in many *Leishmania* sp., those of the subgenus *Viannia* retained a functional pathway, opening up its use as a tool for genetic analysis. In this work we describe a useful Gateway™ site specific recombination based system for rapidly and efficiently generating stem-loop constructs suitable for RNAi tests, and applied it towards a spectrum of *Leishmania* genes of interest to illustrate its range and potential. For perspective, **Table 2** summarizes the genes and the outcomes obtained in this or previous studies.

**Table 2.** Summary of genes targeted by RNAi in *L. (Viannia) sp.* and their phenotypes.

| Gene | Gene ID | Results | Reference |
| --- | --- | --- | --- |
| *LbrAGO1* | LbrM.11.0360 | Reduced RNAi activity | [10] |
| *LbrLPG1* | LbrM.25.0010 | Little mRNA change; remains LPG+ | [10] |
| *LbrLPG2* | LbrM.20.2700 | 3-fold lower mRNA; remains LPG+ | [10] |
| *LbrLPG3* | LbrM.29.0780 | 3-fold lower mRNA; remains LPG+ | [10] |
| *LbrHGPRT* | LbrM.21.0990 | Reduced protein | This work |
| *LbrPFR1* | LbrM.31.0160 | Reduced mRNA; abnormal swimming | [10] |
| *LbrPFR2* | LbrM.16.1480 | Reduced mRNA; abnormal swimming | [10] |
| *Leishmania*<br>RNA virus 1 | *LgyM4147*<br>*LbrLEM2700*<br>*LbrLEM2780*<br>*LbrLEM3874* | LRV1 elimination | [19] |
| *Lbr δ*<br>Amastin | Lbr.M20.0160<br>(gene family) | Reduced mRNA; impaired viability of<br>intracellular amastigotes | [21] |
| *LgyVTC4* | Lgy4147 VTC4 | Reduced mRNA; polyphosphate levels<br>decreased; little impact on growth | [53] |
| *LbrAPRT1* | LbrM.26.0130 | Resistant to APP | This work |
| *LbrQDPR* | LbrM.20.3970 | Altered drug sensitivity | This work |
| *LbrA600* | LbrM.20.3230 | Growth defect as amastigotes | This work |
| *LbrIFT140*<br>(retrograde) | LbrM.32.0380 | Unable to recover transfectants of WT *Lbr* | This work |
| *LbrIFT122*<br>(retrograde) | LbrM.35.1120 | Unable to recover transfectants from WT *Lbr* | This work |
| *LbrIFT172*<br>(anterograde) | LbrM.21.1210 | Unable to recover transfectants from WT *Lbr* | This work |
| *Lbr BBS1*,<br>*BBS3*, *BBS4* | LbrM.34.4180<br>LbrM.16.1430<br>LbrM.35.2520 | No effect on growth or morphology | This work (data not<br>shown) |

Of the 22 targets listed, 6 (27%) showed changes in survival, growth and/or morphology when knocked down (*PFR1, PFR2, IFT140, IFT122, IFT172, VTC4*). When other criteria were evaluated phenotypes were detected in a further 9 (41%; *APRT1*, *A600*, *QDPR*, *AGO1*, δ-amastin, and 4 LRV1s). In contrast, no phenotypes were detected with the methods applied with 7 target genes (*HGPRT1*, *LPG1*, *LPG2*, *LPG3*, *BBS1*, *BBS3* or *BBS4*). Our findings can be compared with the more extensive studies from African trypanosomes, where about 1/3 of genes targeted in chromosomal surveys or genome wide studies show changes in survival, growth or morphology [16,54]. As in trypanosomes, genes involved in the synthesis of abundant surface glycoconjugates such as *Leishmania* lipophosphoglycan *LPG1-3* showed little phenotype, while other genes such as those implicated in the parasite flagellum (*PFR* or *IFT*) yielded readily detected phenotypes. Importantly phenotypes were obtained in several repetitive gene families, including *Lbr A600*, *PFR*s, δ*-amastins* and *QDPR*, the latter posing an addition complexity as an ‘interspersed’ repetitive gene family. Similarly we have been able to target cytoplasmic RNA viruses effectively such as the LRV1 totiviruses of *Lbr* and *Lgy*. These data suggest that RNAi offers another strong option for functional genomics in *Leishmania*.

The specificity of RNAi knockdowns can be be assessed in several ways. First, the dependency of RNAi on a functional Argonaute (*AGO1*) can be used to support conclusions about the essentiality of a given RNAi target, as illustrated by studies with several essential *IFT* genes (**Table 1**). Secondly, the highly expressed integrated SSU:IR-StL constructs used here typically yield very high levels of siRNAs and occasionally dsRNA, both of which can have off target effects [19]. The specificity for the target gene may be assessed by reintroduction of a ‘recoded’ RNAi-resistant target gene [21], or by selection for rare spontaneous excision of the integrated SSU:IR-StL construct from the rRNA gene array [55], both of which should restore the WT phenotype.

While most studies summarized in **Table 2** have been carried out with *L. braziliensis*, RNAi was also effective in *L. guyanensis* (*VTC4, LRV1*). However, in previous work we showed in both LUC and LRV1 studies that the efficacy of RNAi was significant less than in *Lbr* [10,19]. The factors responsible have not been studied, and in our previous studies we noted that even with strong RNAi induction target RNA levels sometimes did not decline [10]. Thus, the ‘penetrance’ of RNAi efficacy can vary widely amongst specific genes, and/or species or strains, something that should be considered in future studies. Fortunately, the ease by which RNAi constructs may be generated using site-specific recombinase technology (**Figure 1**) allows this to be explored with minimal effort.

Interestingly, with the selectable *APRT1*/APP system we were able to increase the strength of RNAi nearly 7 fold in *Lbr* (**Figure 7**), suggesting it may be possible in the future to develop lines where the efficacy of RNAi is enhanced. The feasibility of this was shown in mammalian cells where increased expression of Argonaute-2 enhanced RNAi activity [26].

While the introduction of CRISPR/Cas9 technology provides an attractive alternative to RNAi for gene ablation, there are some useful applications of RNAi as well. Inducible RNAi systems are readily reversible, and the stem length dependency shown here offers the possibility of developing ‘graded’ RNAi responses to yield stable hypomorphic lines. This was illustrated with the *Lbr IFT140* gene, where shorter stems led to viable cells, and by varying stem length, ones could be found yielding viable cells (at a reduced frequency) showing flagellar defects (**Figure 5**), allowing further exploration of the impact of RNAi on flagellar biology or cell physiology.

## Supplementary Materials

The following supporting information can be downloaded at: *JOURNAL WILL PROVIDE LINK:*

**Supplemental Table S1**: Molecular constructs.
**Supplemental Table S2**: Oligonucleotide primers.
**Supplemental Figure S1**: Western blot of *HGPRT*-StL transfectants. This figure shows the western blot whose quantitation is shown in **Figure 2A**.
**Supplemental File S1**: Sequence of the pIR1*HYG*-GW vector.

## Author Contributions

Study conception: L-FL., SMB.; Experimental design, performance, data analysis: L.-F.L., K.L.O., S.J., J.E.M, E.A.B and SMB; Manuscript preparation: S.M.B. and L.-F.L.; Supervision and funding acquisition: S.M.B. All authors have read and agreed to inclusion as authors of this manuscript.

## Funding

This work was supported by NIH grant R01 AI029646 (SMB).

## Institutional Review Board Statement

Not applicable (no vertebrate animals).

## Informed Consent Statement

Not applicable.

## Data Availability Statement

All data are included in the text or supplementary material.

## Acknowledgments

We thank D. McMahon-Pratt and S.C. Alfieri for providing strains of M2903 *Leishmania*, J. Boitz and B. Ullman (Oregon Health Sciences University) for the antisera against *Leishmania* HGPRT and APRT, A. Fairlamb (University of Dundee) for anti-QDPR antiserum, Wandy Beatty for carrying out EM studies, and S.M.F. Murta (WUSM) for assistance with generation of Gateway StL constructs, and Deborah Dobson (WUSM) for comments on this manuscript.

## Conflicts of Interest

The authors declare no conflict of interest.

---

**Supplemental Figure S1.**
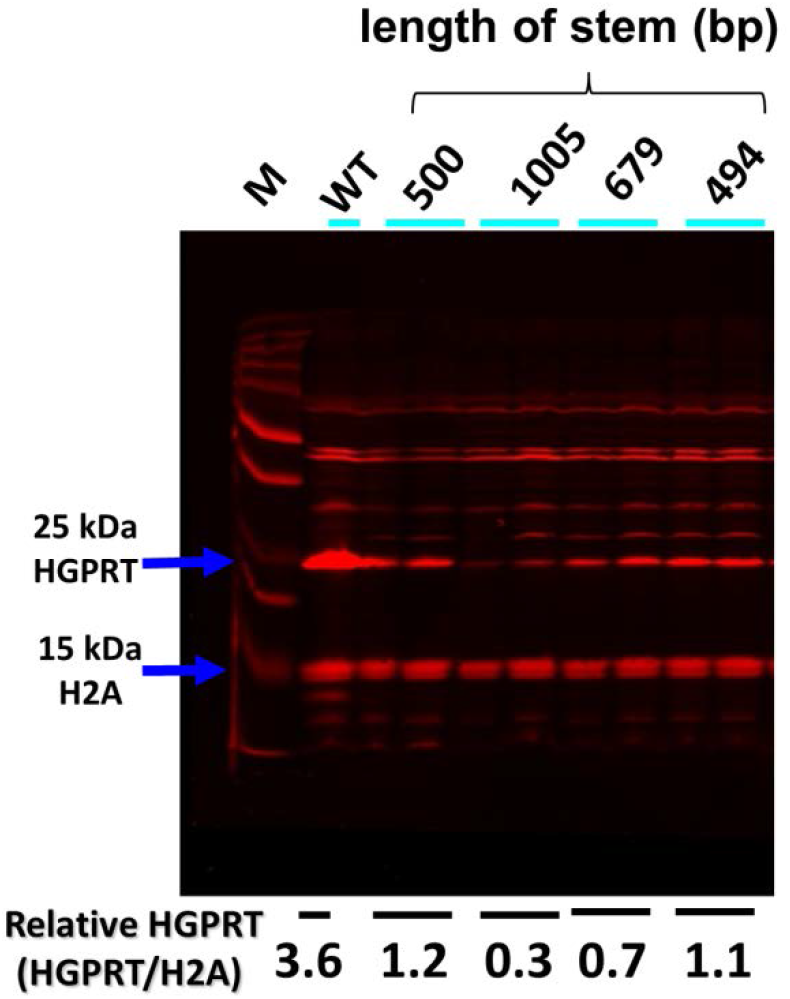
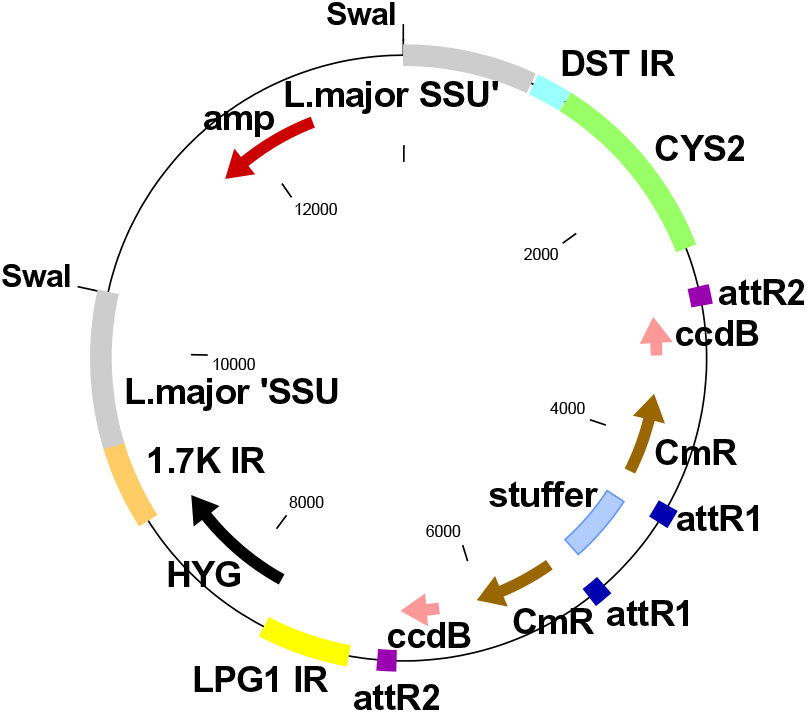
Western Blot analysis of *HGPRT*-StL transfectants. *HGPRT* StL constructs with varying stem lengths (500, 1005, 679, 499 bp) were transfected into *L. braziliensis* and clonal lines recovered. Two of each were subjected to Western blotting with anti-HGPRT and anti-H2A, and the relative expression was calculated and normalized to WT. M, molecular weight marker; WT, *Lbr*.

**Supplemental File 1. Sequence of a representative pIR-GW vector, pIR1HYG-GW**

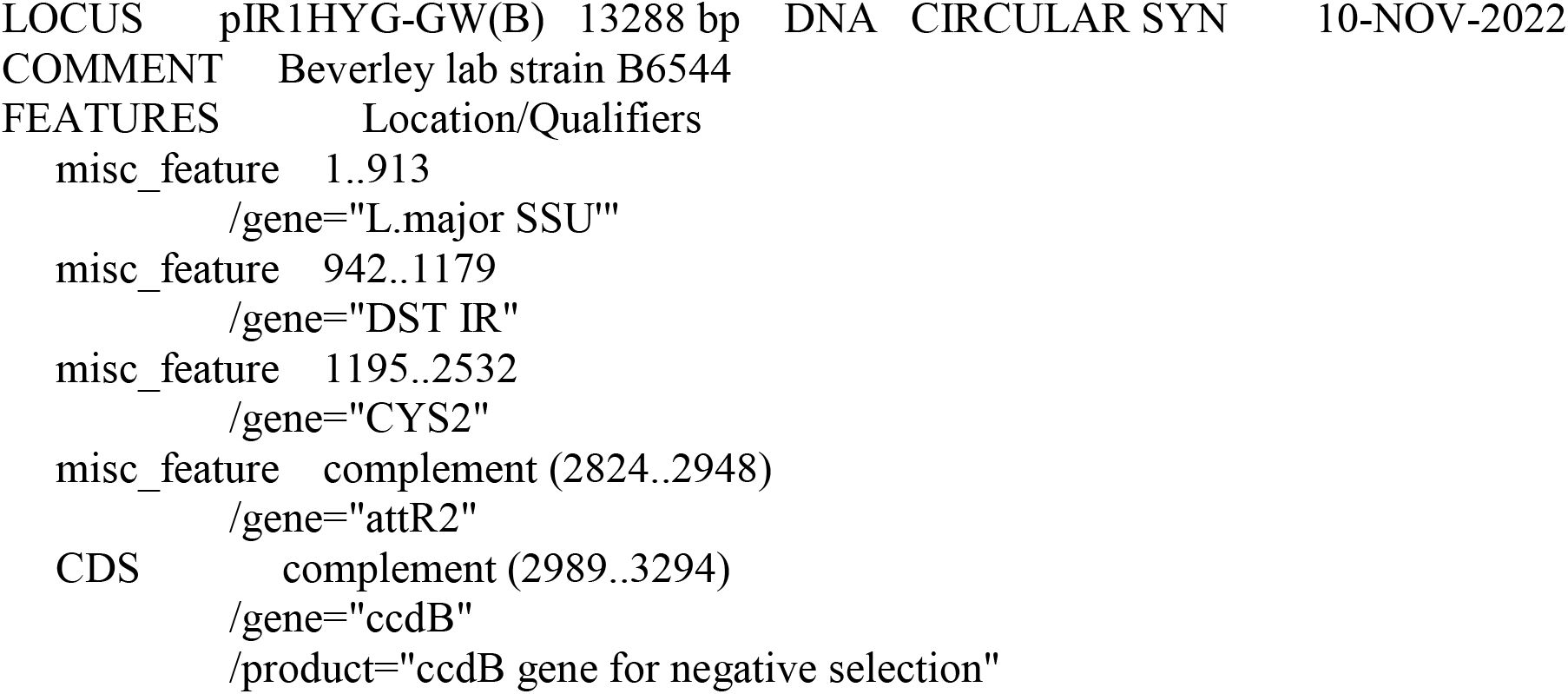
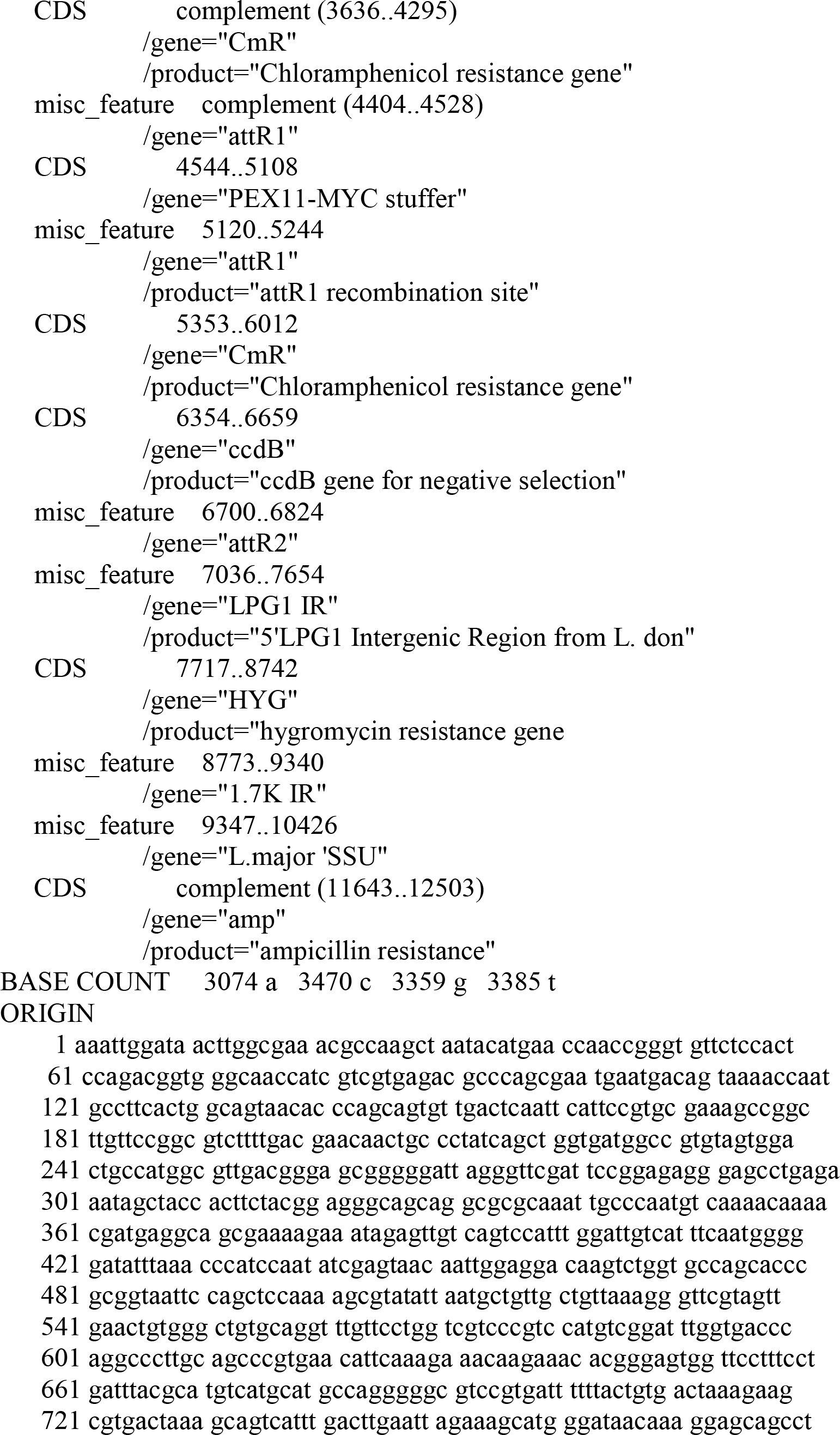
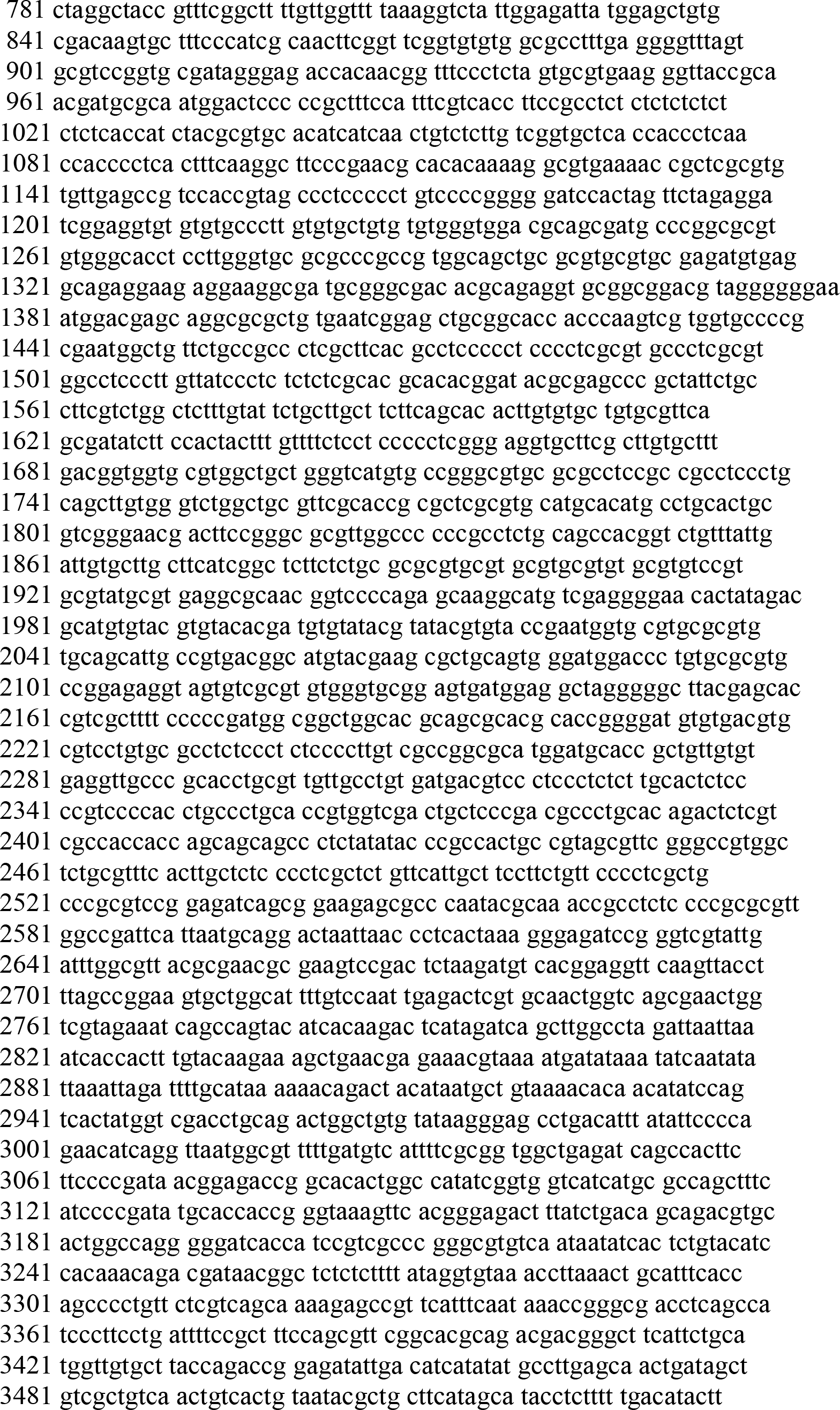
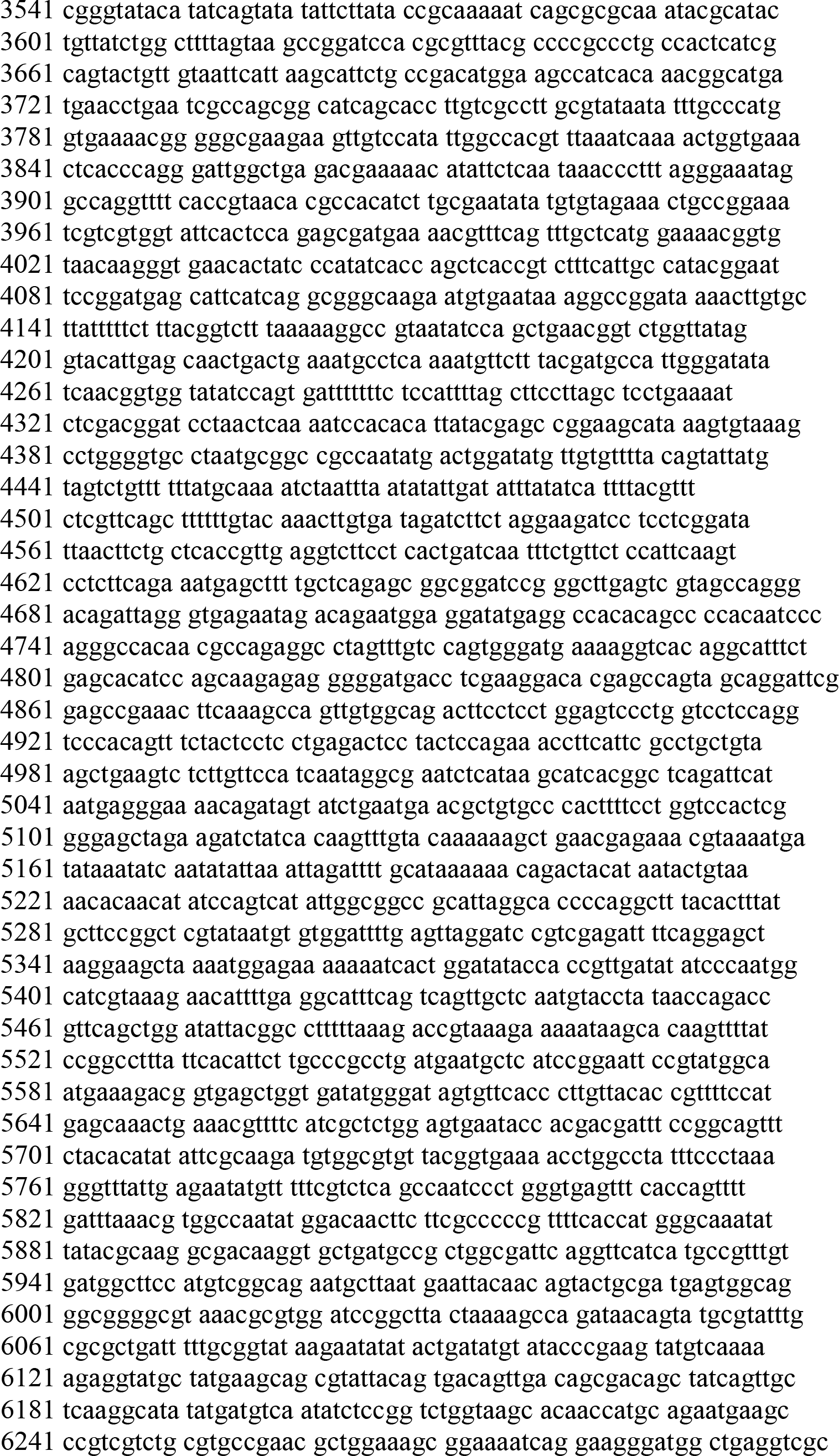
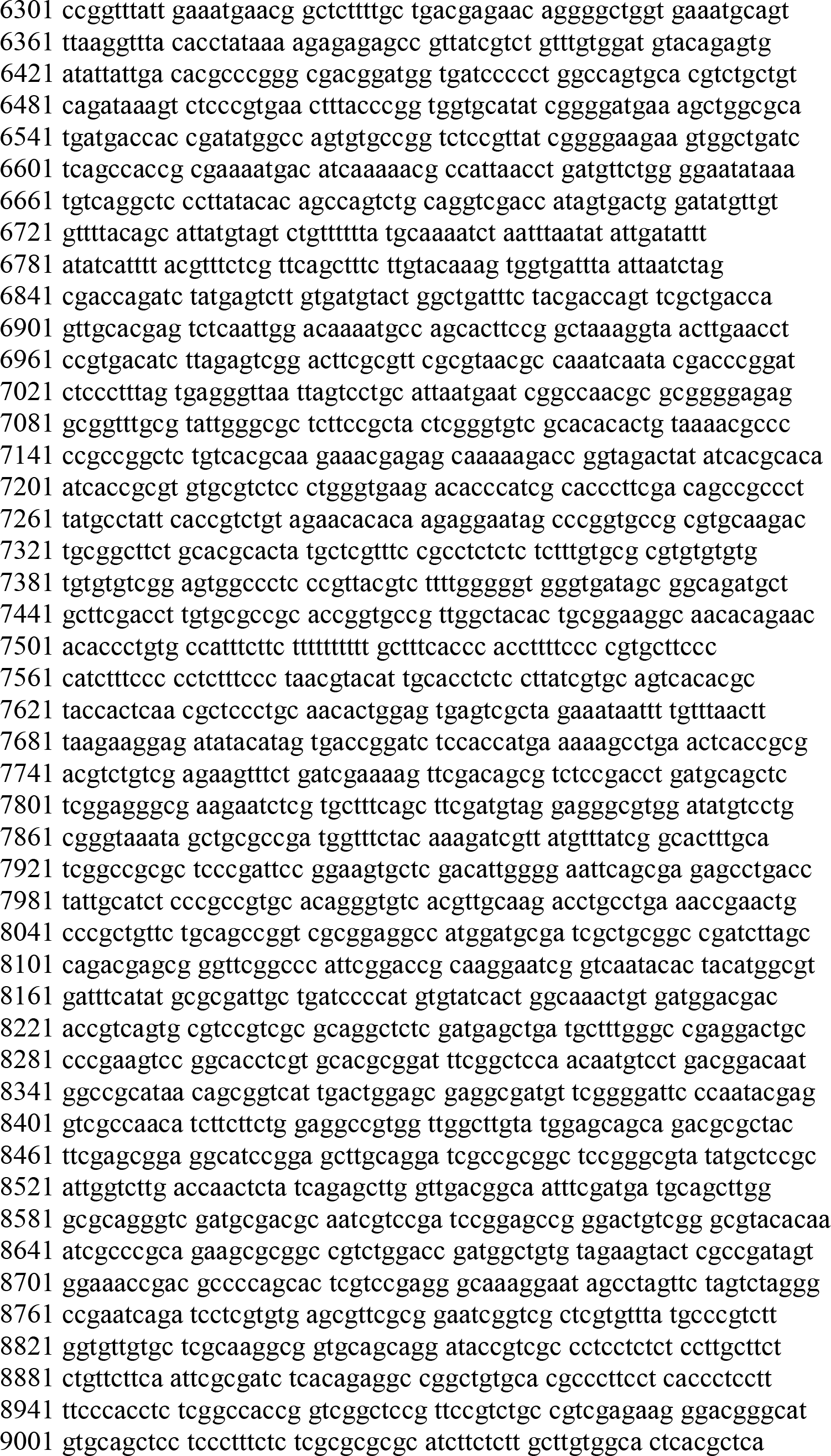
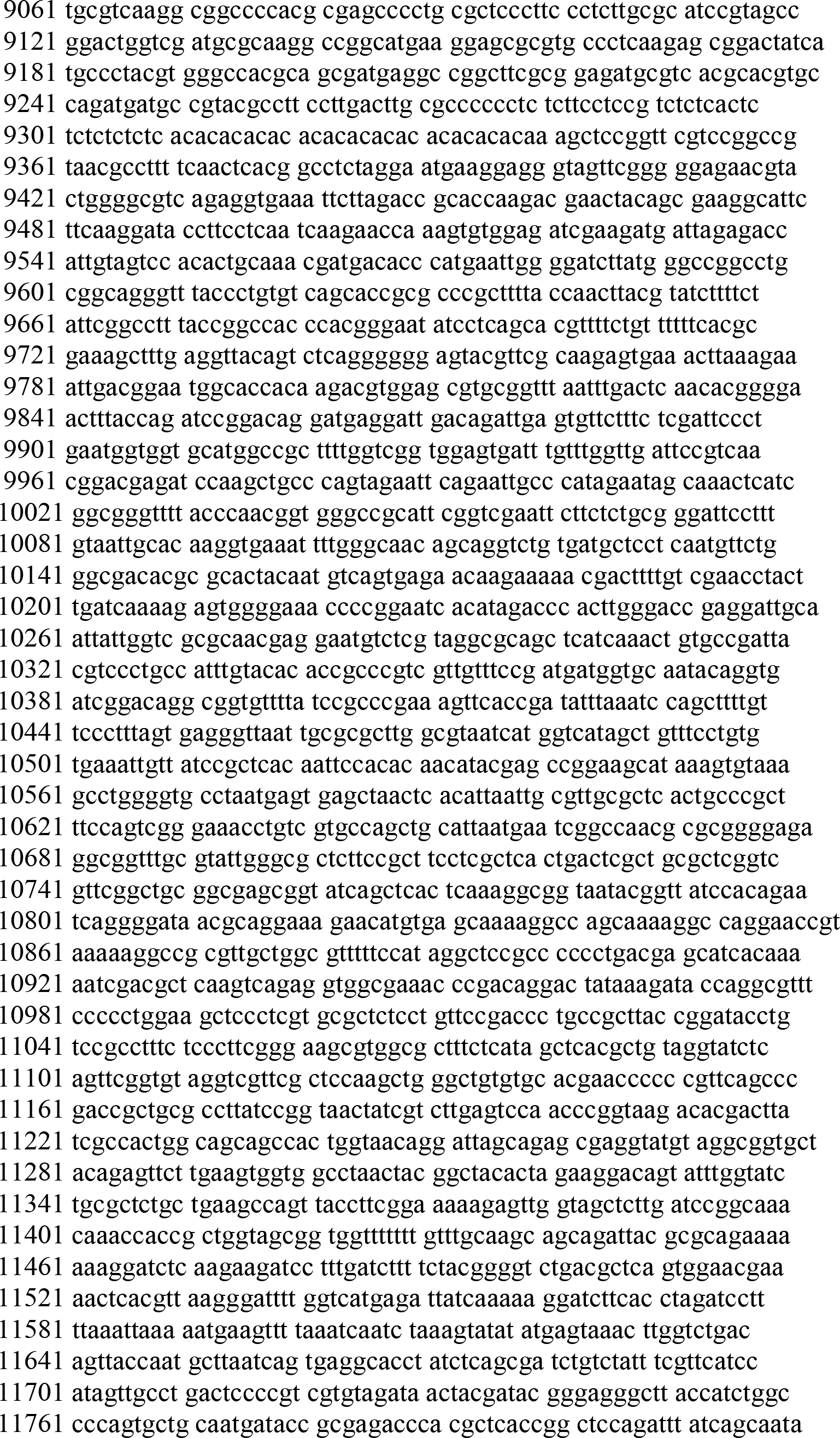
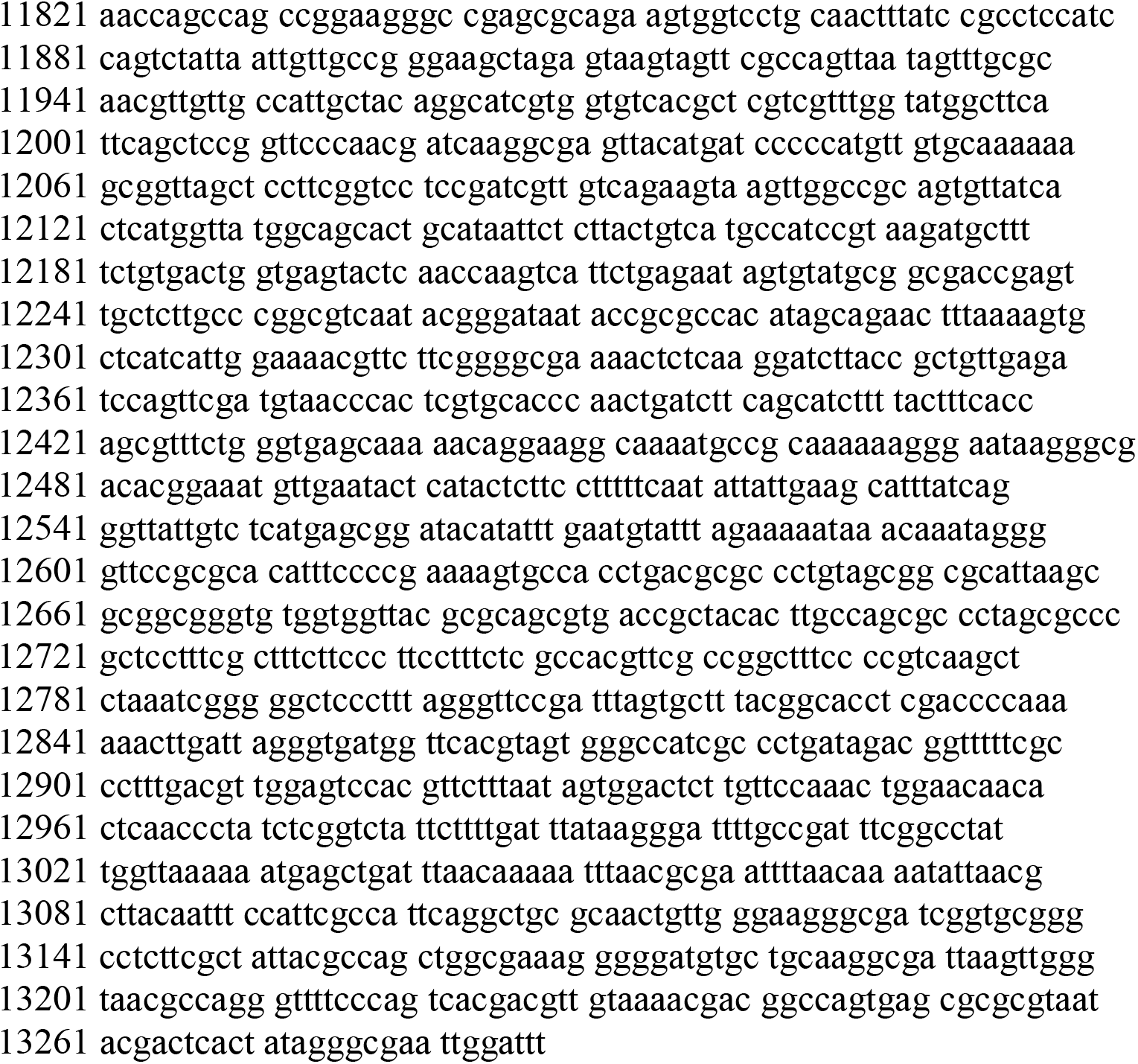

**Table S1.** Molecular constructs used. (oligonucleotide primers can be found in Supporting Information Table S2)

| Constructs | Lab strain number | Description or method of construction |
| --- | --- | --- |
| pGEM-T Easy | commercial | *E. coli* expression vector; Amp<sup>R</sup> (Promega; Catalog numbe: A137A) |
| pCR™8/GW/TOPO™ | commercial | TA Cloning vector for creating entry clone for Gateway™ recombination cloning; Spectinomycin<sup>R</sup> (Thermo Fisher Scientific; Catalog number: K252020) |
| pGEMT- stuffer | B5974 | *Bgl*II-*Nhe*I-PEX11-AvrII-*Bgl*I fragments were amplified from pIR1SAT-GFP65-StL (B4733) with primers SMB2817/2818 then cloned into pGEM-T Easy; Amp<sup>R</sup>. Lye *et al* (2010) *PLoS Pathogens* 6: e1001161 |
| pGEMT-stuffer -one GW | B6158 | The Gateway® cassette (GW) contains *ccdB*, a lethal gene that targets DNA gyrase, and *CmR* encoding chloramphenicol-resistance, flanked by two *attR* sequences necessary for site-specific recombination (*attR1-ccdB-CmR-attR2*) was amplified from pDONR221 (Invitrogen) with primers S1/S2, then cloned into the *Nhe*I site of the pGEMT-stuffer (B5974). |
| pGEMT-stuffer - 2 GW divergent | B6218 | The Gateway® cassette fragment (*attR1-ccdB-CmR-attR2*) was amplified from pDONOR221 (Invitrogen) with primers described in the methods, and inserted by blunt end ligation into the *Avr*II site of the pGEMT-stuffer one GW (B6158) in the divergent orientation. |
| pIR1SAT-GW | B6223 | The *Sph*I/*Hin*dIII fragment containing the divergent stuffer/ GW cassettes was released from pGEMT-stuffer- 2 GW divergent (B6218) and blunt end ligated into the *Sma*I site of pIR1SAT |
| pIR2SAT | B6351 | Lye et al PMID: PMC2965760 PMID: 21060810 *L.braziliensis* α-tubulin IG was amplified with SMB3627 and 3635 and cloned into the *Xma*I and *Bgl*II site of pIR1SAT (removed *Xma*I-*Bgl*II *Lp* Cyst2 IR) |
| pIR2HYG | B6441 | Lye et al PMID: PMC2965760 PMID: 21060810 The *Bam*HI-CCACC-HYG-*Spe*I fragment was amplified with pXGHYG (B3318) and SMB3828/3829 then digested with *Bam*HI/*Spe*I and swapped into *Bam*HI/*Spe*I digested of pIR2SAT (B6351) |
| pIR1SAT | B3541 | Robinson *et al* (2003) *Mol Biochem Parasitol* 128: 217–22 |
| pIR1HYG | B6174 | The *HYG* ORF wa amplified from pXGHYG (B3318) with primers SMB2819/2820 adding flanking *Bgl*II and *Sly*I sites and ligated into the *Bam*HI/*Spe*I backbone of pBS-IR-HYG (B6048) |
| pIR1PAC | B6176 | The *PAC* ORF was amplified from pXGPAC (B3325) with primers SMB2821/2822 adding flanking *Bgl*II and *Sly*I sites and ligated into the *Bam*HI/*Spe*I backbone of pBS-IR-PAC (B6060) |
| pIR1BSD | B6173 | The *BSD* ORF was amplified from pXGBSD (B4098) with primers SMB2823/2824 adding flanking *Bgl*II and *Sly*I sites and ligated into the *Bam*HI/*Spe*I backbone of pBS-IR-BSD (B6052) |
| pXG1a | B1495 | pXG1A was created by partial digestion and fill in of the *Spe*i site of the expression site of pXG (B1288; Ha et al (1996; *Mol. Biochem. Parasitol.* 77:57-64) |
| pXGHYG | B3318 | A 1.05 kb *HYG* ORF cassette from pXG63HYG (B617) ligated to 5.9kb fragment of pXG1a/*Spe*I. |
| pXGPAC | B3325 | A 720 bp *PAC* ORF cassette was ligated to 5.9kb fragment of pXG1a/*Spe*I |
| pXGBSD | B4098 | Guo *et al* (2017), *J. Biol. Chem.* 292: 10696-10708. |
| pIR1Phleo -GFP65*- LUC | B6034 | Lye *et al* (2010) *PLoS Pathogens* 6: e1001161 |
| pIR1SAT-LUC-StL | B6190 | Lye *et al* (2010) *PLoS Pathogens* 6: e1001161 |
| pIR2SAT-LUC (B) | B6367 | Firefly luciferase (LUC) ORF was released from pIR1Phleo -GFP65*- LUC (B6034) by *Bgl*II digestion an inserted and cloned into the B site of *Leishmania* expression vector pIR2SAT |
| pIR2SAT-LUC-StL(A)-LUC (B) | B6386 | *Bgl*III LUC-StL fragment was released from pIR1SAT-LUC-StL (B6190) and filled in then cloned into *Sma*I site (A site) of pIR2SAT-LUC (B) (B6367) |
| pIR2SAT-LUC-StL(A)-*APRT1*-StL (B) | B6391 | *APRT1*-StL fragment was released from pIR1*BSD*-*APRT1*-StL (B6742) and replaced the LUC ORF in the B site of the pIR2SAT-LUC-StL(A)-LUC (B) (B6386) |
| pIR1*HYG*-GW | B6544 | The *AhdI*/*Sap*I fragment bearing the inverted *ccdB-CmR* / Loop from pIR1SAT-GW (B6223) was blunt-end ligated into the *Bgl*II site of pIR1*HYG* (B6174) |
| pIR1*PAC*-GW | B6543 | The *AhdI*/*Sap*I fragment bearing the inverted *ccdB-CmR* / Loop from pIR1SAT-GW (B6223) was blunt-end ligated into the *Bgl*II site of pIR1*PAC* (B6176) |
| pIR1*BSD*-GW | B6542 | The *AhdI*/*Sap*I fragment bearing the inverted *ccdB-CmR* / Loop from from pIR1SAT-GW (B6223) was blunt-end ligated into the *Bgl*II site of pIR1*BSD* (B6173) |
| pIR2*HYG*-GW | B6563 | The *AhdI*/*Sap*I fragment bearing the inverted *ccdB-CmR* / Loop from pIR1SAT-GW (B6223) was blunt-end ligated into the *Bgl*II site of pIR2*HYG* (B6441). Brettmann et al (2016), Proc. Natl. Acad. Sci. USA 113:11998-12005 |
| pIR2SAT-GW | B6566 | The *AhdI*/*Sap*I fragment bearing the inverted *ccdB-CmR* / Loop from pIR1SAT-GW (B6223) was blunt-end ligated into the *Bgl*II site of pIR2SAT (B6351). Brettmann et al (2016), Proc. Natl. Acad. Sci. USA 113:11998-12005 |
| **<u>STL constructs</u>** |  |  |
| pIR2*HYG*-*LbrA600*-StL (519) | B6595 | A 519 nt stem from *LbrA600* (LbrM.26.0130) was obtained by PCR with primers SMB 4144/4145, cloned into pCR8/GW/TOPO and used as donor with pIR2*HYG*-GW |
| pIR2SAT-*LbrIFT140*-StL (940) | B6666 | A 940 nt stem of *LbrIFT140* (LbrM.32.0380) was obtained by PCR with primers SMB 4179/4180 , cloned into pCR8/GW/TOPO and used as donor with pIR2*HYG*-GW |
| pIR2SAT-*LbrIFT122*-StL (1100) | B6664 | A 1100 nt stem of *LbrIFT122* (LbrM.35.1120) obtained by PCR with primers SMB 4181/4182), cloned into pCR8/GW/TOPO and used as donor with pIR2*HYG*-GW |
| pIR2SAT-*LbrIFT172*-StL (881) | B6674 | The stem (881 nt) of *LbrIFT172* (LbrM.21.1210) obtained by PCR with primers SMB 4175/4176, cloned into pCR8/GW/TOPO and used as donor with pIR2SAT-GW |
| pIR1SAT-*LbrQDPR*-StL (588) | B6296 | The stem (588 nt) of *LbrQDPR* (LbrM.20.3970) obtained by PCR with primers SMB 3549/3550, cloned into pCR8/GW/TOPO and used as donor with pIR1SAT-GW |
| pIR1*BSD*-*LbrAPRT1*-StL (514) | B6742 | The stem (514 nt) of *LbrAPRT1* (LbrM.26.0130) obtained by PCR with primers SMB 3460/3461, cloned into pCR8/GW/TOPO and used as donor with pIR1*BSD*-GW |
| **<u>HGPRT deletion series STL constructs</u>** |  |  |
| pIR1SAT-*LbrHGPRT*-ORF alone-StL (500) | B6136 | The stem (500 nt) of *LbrHGPRT* (LbrM.21.0990) obtained by PCR with primers SMB 2902/2903, cloned into pCR8/GW/TOPO and used as donor with pIR1SAT-GW |
| pIR1SAT-*LbrHGPRT*-3' alone-StL (494) | B6234 | The stem (494 nt) of *LbrHGPRT* (LbrM.21.0990) obtained by PCR with primers SMB 3489/3468, cloned into pCR8/GW/TOPO and used as donor with pIR1SAT-GW |
| pIR1SAT-*LbrHGPRT*-5'+ORF-StL (679) | B6241 | The stem (679 nt) of *LbrHGPRT* (LbrM.21.0990) obtained by PCR with primers SMB 3487/2903, cloned into pCR8/GW/TOPO and used as donor with pIR1SAT-GW |
| pIR1SAT-*LbrHGPRT*-ORF+3'-StL (1005) | B6243 | The stem (1005 nt) of *LbrHGPRT* (LbrM.21.0990) obtained by PCR with primers SMB 2902/23468, cloned into pCR8/GW/TOPO and used as donor with pIR1SAT-GW |
| **<u>IFT140 deletion series-StL constructs</u>** |  |  |
| pIR2SAT-*LbrIFT140*-StL ( 941) | B6666 | The stem (941 nt) of *LbrIFT140* (LbrM32.0380) obtained by PCR with primers SMB 2902/2903, cloned into pCR8/GW/TOPO and used as donor with pIR2SAT-GW |
| pIR2SAT-*LbrIFT140*-StL (562) | B6857 | The stem (562 nt) of *LbrIFT140* (LbrM32.0380) obtained by PCR with primers SMB 4675/4180, cloned into pCR8/GW/TOPO and used as donor with pIR2SAT-GW |
| pIR2SAT-*LbrIFT140*-StL (497) | B6954 | The stem (497 nt) of *LbrIFT140* (LbrM32.0380) obtained by PCR with primers SMB 4964/4180, cloned into pCR8/GW/TOPO and used as donor with pIR2SAT-GW |
| pIR2SAT-*LbrIFT140*-StL (431) | B6953 | The stem (431 nt) of *LbrIFT140* (LbrM32.0380) obtained by PCR with primers SMB 4965/4180, cloned into pCR8/GW/TOPO and used as donor with pIR2SAT-GW |
| pIR2SAT-*LbrIFT140*-StL (365) | B6952 | The stem (365 nt) of *LbrIFT140* (LbrM32.0380) obtained by PCR with primers SMB 4966/4180, cloned into pCR8/GW/TOPO and used as donor with pIR2SAT-GW |
| pIR2SAT-*LbrIFT140*-StL (298) | B6858 | The stem (298nt) of *LbrIFT140* (LbrM32.0380) obtained by PCR with primers SMB 4676/4180, cloned into pCR8/GW/TOPO and used as donor with pIR2SAT-GW |
| pIR2SAT-*LbrIFT140*-StL (131) | B6891 | The stem (131nt) of *LbrIFT140* (LbrM32.0380) obtained by PCR with primers SMB 4677/4180, cloned into pCR8/GW/TOPO and used as donor with pIR2SAT-GW |
| **<u>LUC deletion series StL constructs</u>** |  |  |
| pIR2HYG-*LUC*-StL (940) | B6896 | The stem (940 nt) of firefly luciferase (*Photinus pyralis*) obtained by PCR with primers SMB 4818/4817, cloned into pCR8/GW/TOPO and used as donor with pIR2HYG-GW |
| pIR2HYG-*LUC*-StL (533) | B6897 | The stem (533 nt) of firefly luciferase (*Photinus pyralis*) obtained by PCR with primers SMB 4819/4817, cloned into pCR8/GW/TOPO and used as donor with pIR2HYG-GW |
| pIR2HYG-*LUC*-StL (285) | B6898 | The stem (285 nt) of firefly luciferase (*Photinus pyralis*) obtained by PCR with primers SMB 4820/4817, cloned into pCR8/GW/TOPO and used as donor with pIR2HYG-GW |
| pIR1BSD-*LUC*-StL (158) | B6959 | The stem (158 nt) of firefly luciferase (*Photinus pyralis*) obtained by PCR with primers SMB 4957/4958, cloned into pCR8/GW/TOPO and used as donor with pIR1BSD-GW |
| pIR2HYG-*LUC*-StL (128) | B6899 | The stem (128 nt) of firefly luciferase (*Photinus pyralis*) obtained by PCR with primers SMB 4821/4817, cloned into pCR8/GW/TOPO and used as donor with pIR2HYG-GW |
| pIR2HYG-*LUC*-StL (54) | B6900 | The stem (54 nt) of firefly luciferase (*Photinus pyralis*) obtained by PCR with primers SMB 3487/2903, cloned into pCR8/GW/TOPO and used as donor with pIR2HYG-GW |

**Table S2.**
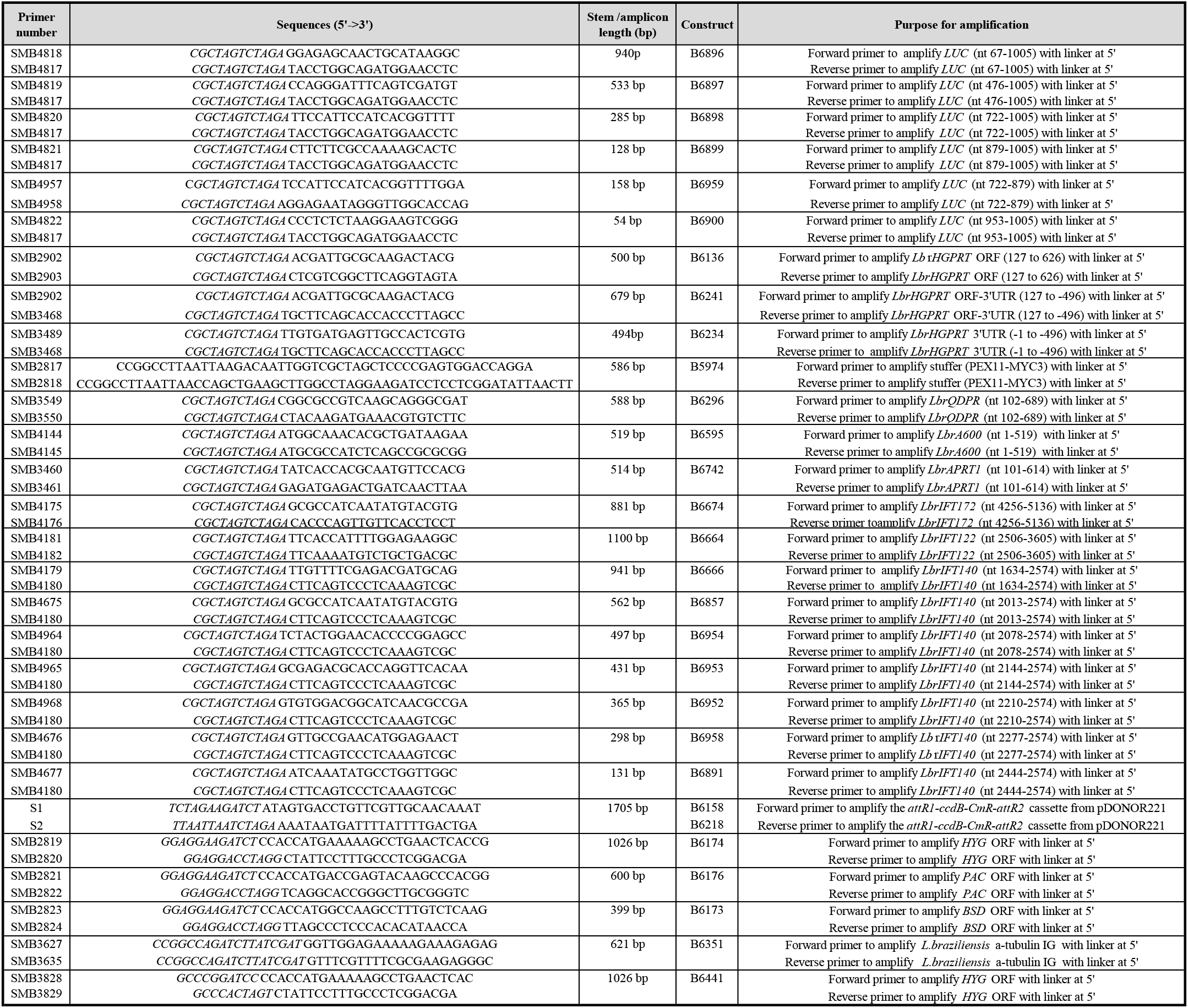
Oligonucleotide primers.

## Notes

### Competing Interest Statement

The authors have declared no competing interest.

